# The ESX-1 substrate PPE68 has a key function in the ESX-1 mediated secretion in *Mycobacterium marinum*

**DOI:** 10.1101/2022.06.21.497121

**Authors:** Merel P.M. Damen, Aniek S. Meijers, Esther M. Keizer, Sander R. Piersma, Connie R. Jiménez, Coenraad P. Kuijl, Wilbert Bitter, Edith N. G. Houben

## Abstract

Mycobacteria use specialized type VII secretion systems (T7SSs) to secrete proteins across their diderm cell envelope. One of the T7SS subtypes, named ESX-1, is a major virulence determinant in pathogenic species such as *Mycobacterium tuberculosis* and the fish pathogen *Mycobacterium marinum.* ESX-1 secretes a variety of substrates, called Esx, PE, PPE and Esp proteins, at least some of which as folded heterodimers. Investigations into the functions of these substrates is problematic, because of the intricate network of co-dependent secretion between several ESX-1 substrates. Here, we describe that the ESX-1 substrate PPE68 is essential for secretion of the highly immunogenic substrate EsxA and EspE via the ESX-1 system in *M. marinum*. While secreted PPE68 is processed on the cell surface, the majority of cell-associated PPE68 of *M. marinum* and *M. tuberculosis* is present in a cytosolic complex with its PE partner and the EspG_1_ chaperone. Interfering with the binding of EspG_1_ to PPE68 blocked its export and the secretion of EsxA and EspE. In contrast, *esxA* is not required for the secretion of PPE68, revealing hierarchy in co-dependent secretion. Remarkably, the final ten residues of PPE68, a negatively charged domain, seem essential for EspE secretion, but not for the secretion of EsxA and PPE68 itself. This indicates that distinctive domains of PPE68 are involved in secretion of the different ESX-1 substrates. Based on these findings, we propose a mechanistic model for the central role of PPE68 in ESX-1 mediated secretion and substrate co-dependence.

**Importance:** *Mycobacterium tuberculosis* use type VII secretion systems (T7SSs) to secrete proteins across its uniquely hydrophobic diderm cell envelope. A T7SS subtype, called ESX-1, is one of its most important virulence determinants by mediating intracellular survival through phagosomal rupture and subsequent translocation of the mycobacterium to the host cytosol. Identifying the ESX-1 substrate that is responsible for this process is problematic because of the intricate network of co-dependent secretion between ESX-1 substrates. Here, we provide mechanistic insight into the central role of the ESX-1 substrate PPE68 for the secretion of ESX-1 substrates, using the model organism *Mycobacterium marinum*. Unravelling the mechanism of co-dependent secretion will aid the functional understanding of T7SSs and will allow the analysis of the individual roles of ESX-1 substrates in the virulence caused by this significant human pathogen.

## Introduction

The genus Mycobacterium belongs to the phylum Actinobacteria and includes the major human pathogens *Mycobacterium tuberculosis* and *Mycobacterium leprae*. Mycobacteria have a distinctive cell envelope that is shared with other bacteria within the order of the Corynebacteriales. The unique feature of their cell envelope are the long chained fatty acids, called mycolic acids, that assemble into a second hydrophobic bilayer, also known as the mycobacterial outer membrane (1). Mycobacteria have acquired specialized secretion systems, called the type VII secretion systems (T7SS), to export proteins across this multi-layered cell envelope. There are five paralogous T7SSs present in *M. tuberculosis*, called ESX-1 to ESX-5. The ESX-1 system is considered a major virulence factor of *M. tuberculosis* and the fish pathogen *Mycobacterium marinum* through its essential role in phagosomal rupture and subsequent translocation of the pathogens to the host cytosol (2–5). As a consequence, the absence of the *esx-1* region results in severe attenuation of pathogenic mycobacteria, which is further illustrated by the observation that a large deletion within the same region (RD1) is the prime cause of the attenuation of the live vaccine strain *Mycobacterium bovis* BCG (6–8).

The ESX-1 system secretes over a dozen different proteins, which all belong to the EsxAB clan (Pfam CL0352) and can be categorized into four distinctive protein families *i.e.* Esx/WxG100, PE, PPE and Esp. The best-studied and highly immunogenic ESX-1 substrates EsxA and EsxB belong to the Esx/WxG100 protein family, which are characterized by a WxG protein motif and generally are 100 amino acids long. PE35 and PPE68 are encoded within the same operon as *esxA* and *esxB* and belong to PE and PPE protein families that share homologous N-terminal domains of approximately 110 amino acids and 180 amino acids, respectively, with conserved proline–glutamic acid (PE) and proline–proline–glutamic acid (PPE) motifs (9). While each T7SS secretes its own Esx proteins and often also PE and PPE substrates, the Esp proteins seem to be exclusively secreted by the ESX-1 system.

An intriguing feature of the mycobacterial T7SS substrates is that these proteins are secreted as folded heterodimers that are formed in the cytosol (10–13). While these heterodimers are formed by either Esx substrates or a PE protein with a specific PPE protein, available crystal structures reveal highly conserved features. Each complex forms a four-helix bundle, composed of two helix–turn–helix structures in an antiparallel orientation, with a conserved secretion motif (YxxxD/E) extending from the double-helix of one partner protein (14). In addition to the helix–turn–helix, the N-terminal domain of PPE proteins contain an extra region that is relatively hydrophobic and extends from the PE-PPE interface (15). This so-called helical tip is recognized by a cytosolic chaperone, called EspG, in an ESX system-specific manner, which recognition is required for secretion of the PE/PPE pair (11, 12, 14). In addition to the conserved dimeric structure formed by the N-terminal PE and PPE domains, which is thought to be involved in substrate recognition and secretion of the heterodimers, many PE and PPE proteins have highly variable C-terminal domains that could potentially execute specific functions (16–18). Only limited structural information is available for the Esp substrates, although structure predictions indicate that most of these substrates form similar heterodimers as PE/PPE proteins (19–21).

EsxA has been indicated to play a central role in ESX-1 mediated virulence, including the perturbation of phagosomal membranes (2–4, 22–24). However, while *in vitro* biochemical analyses suggest that EsxA has membranolytic activity (22, 25–30), later reports have ascribed at least some of these observations to detergent contaminations during the purification of EsxA (29, 31). Another complicating factor is that mutations in *esxA* abolishes the secretion of other ESX-1 substrates, such as EspA, EspF, EspJ, EspK and EspB in *M. tuberculosis* and/or *M. marinum* (32–34). In numerous cases, the co-dependence for secretion is mutual, as the deletion of specific *esp* genes also results in abolished EsxA secretion (32, 35–38). This complicates the analysis and interpretation of the role of individual ESX-1 substrates in virulence.

Co-dependent secretion has also been observed for substrates of other ESX systems (39–41). The underlying reason for this phenomenon however remains unclear. Previously, we have provided the first mechanistic insight into this process by the successful redirection of PE35_1/PPE68_1, the *M. marinum* paralogues of *esx-1* locus encoded PE35/PPE68, from the ESX-1 to the ESX-5 system (42, 43). Surprisingly, the redirection of these two ESX-1 substrates also resulted in redirection of EsxA_1. This finding suggests a central role of the PE-PPE-EspG complex in the secretion of Esx substrates. Interestingly, deletion of *espG_1_* results in a general secretion defect not only of respective PE/PPE and Esx substrates in *M. marinum*, but also of Esp proteins (24, 33–35, 44, 45), indicating a central role for PE/PPE substrates in the secretion of all known ESX substrate classes. Here, we investigated the role of PPE68 in ESX-1 mediated secretion of EsxA and EspE in *M. marinum* and report that this PPE protein has a central function in the secretion of these two ESX-1 substrates.

## Results

### The *pe35/ppe68* pair is necessary for secretion of ESX-1 substrates EsxA and EspE, and for ESX-1-mediated hemolysis

To study the effect of PPE68 on protein secretion, we created a frameshift mutation *M. marinum* M to minimize polar effects, using the efficient genome editing tool CRISPR1-Cas9 (46). The generated *ppe68* frameshift (fs) mutant that we selected contained a 1 bp insertion at position 181, which results in an early stop codon after amino acid 202 (Figure S1A). As expected, PPE68 could not be detected in pellet or supernatant fraction of this mutant (Figure 1B, lane 2, 7). In the WT strain, we could identify PPE68 in the pellet fraction but not in the supernatant (Figure 1B, lane 7), which is in line with the observations in *M. tuberculosis* (47). The *ppe68* mutant showed a loss of EsxA in the supernatant fraction, while no accumulation of EsxA in the pellet fraction was observed (Figure 1B, lane 2, 7). The lack of accumulation is possibly caused by a WhiB6-controlled negative feedback mechanism resulting in downregulated expression in the presence of an inactive ESX-1 secretion system (48, 49). In agreement, we observed that mRNA levels of *esxA* in the *ppe68* fs mutant was reduced to a comparable level as in an *eccCb_1_* core complex mutant (Figure S1C). Next, we analyzed the secretion of ESX-1 substrate EspE, which is an abundant cell-surface protein of *M. marinum* and can be extracted by the mild detergent Genapol X-080 (50, 51). The *ppe68* fs mutation prevented the secretion of EspE to the cell surface, while the intracellular levels of EspE in the mutant and the WT were comparable (Figure 1B, lane 2, 7). Complementation of the *ppe68* fs mutant with the *pe35/ppe68* pair, expressed from an integrative pMV vector, failed to restore detectable EspE and EsxA secretion (Figure 1B, lane 8); only the introduction of the complete *ppe68* operon, encompassing *pe35*, *ppe68*, *esxB* and *esxA*, using either the pMV or multicopy pSMT3 vector, restored EspE and EsxA secretion (Figure 1B, lane 9-10). Analysis of three times higher amounts of culture supernatant revealed low levels of EsxA secretion in the *ppe68* fs mutant expressing the *pe35/ppe68* pair from a pSMT3 vector, while EspE secretion was fully restored (Figure S1D, lane 4). We have previously shown that overexpression of *esxB_1/esxA_1* in the WT blocked ESX-1 secretion, and that co-overexpression of the *pe35_1/ppe68_1* released this secretion block (42). These data indicate that balanced production of both heterodimers is required for efficient secretion, which could also be true in this situation.

**Figure 1.**
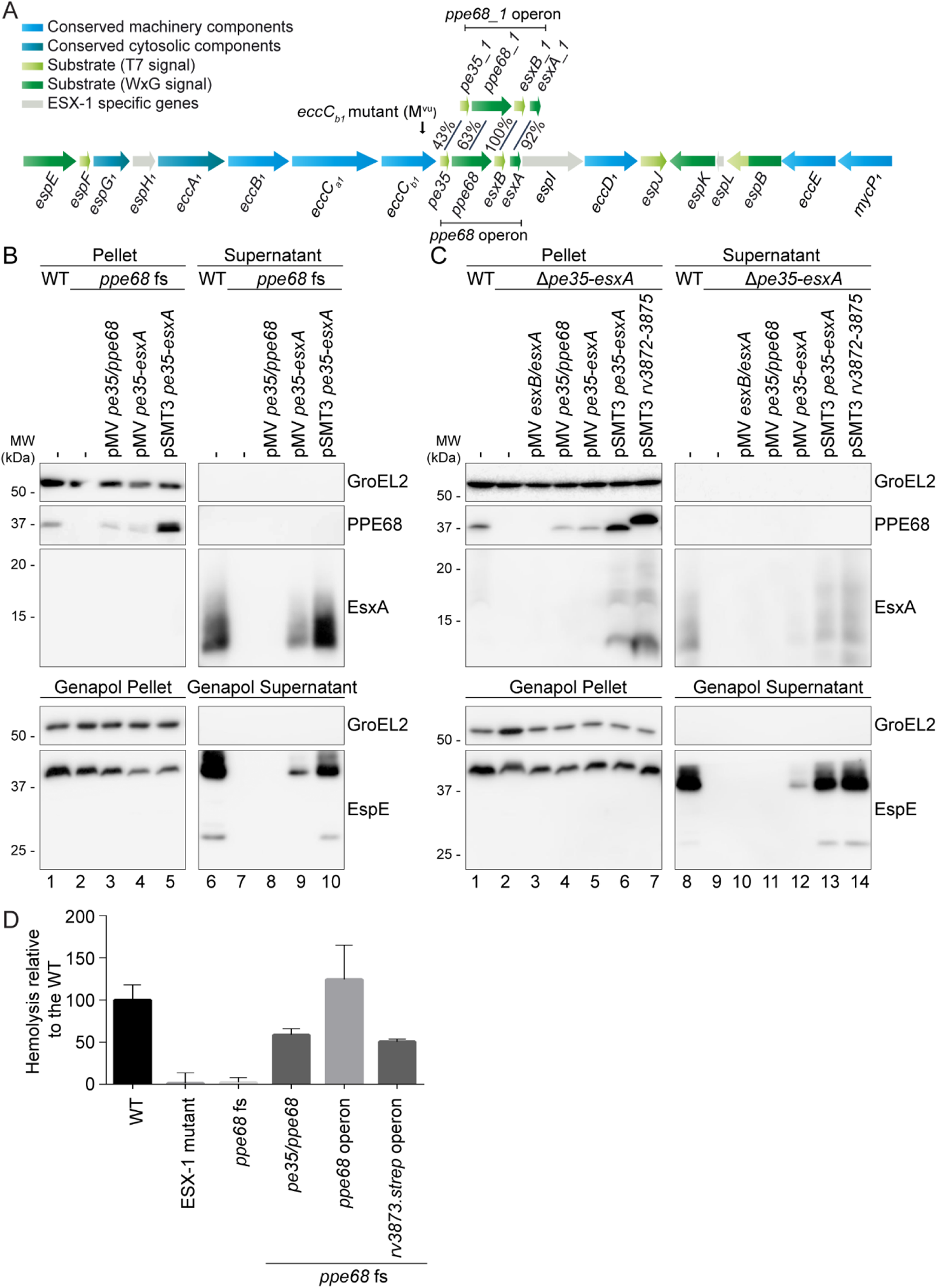
*pe35/ppe68* is necessary for secretion of ESX-1 substrates EsxA and EspE, and is required for ESX-1 mediated hemolysis. A) Schematic overview of the *esx-1* gene cluster and its paralogous region. Genes are color-coded to their subcellular localization (key color). Notably, the *pe* gene upstream of *ppe68_1* was originally named *pe35*, while the paralogous gene upstream of *ppe68* in the *esx-1* locus was named *mmar_5447*. In this study, we renamed these two genes *pe35_1* and *pe35*, respectively. B, C) SDS-PAGE and immunostaining of the cell pellet, culture supernatant, Genapol pellet and Genapol supernatant fractions of *M. marinum* M wild type (WT) and the *ppe68* fs mutant without and with various complementation constructs (B) or WT and the *ppe68* operon mutant (Δ*pe35-esxA*) with and without various complementation constructs (C). Genes were expressed from an integrative pMV vector or a multicopy pSMT3 vector under control of the *hsp60* promotor. In B) the WT, *ppe68* fs mutant and the *ppe68* fs mutant containing a pSMT3 vector also contained a pMV TdTomato plasmid. Proteins were visualized with anti-EsxA, anti-EspE, and anti-PPE68 antibodies (ESX-1 substrates). A processed band of 25 kDa was detected by the anti-EspE antibody in the Genapol supernatant fraction of the WT, which has been reported before (44). As a loading and lysis control, blots were incubated with antibodies directed against the cytosolic GroEL2 protein. In all blots, equivalent OD units were loaded: 0.2 OD for pellet and Genapol pellet and 0.4 OD for supernatant and Genapol supernatant fractions. D) Hemoglobin release after incubation of sheep red blood cells with *M. marinum* WT, an ESX-1 mutant (*eccC_b1_* mutant; M^VU^), the *ppe68* fs mutant, and the *ppe68* fs mutant complemented with either the entire *ppe68* operon of *M. marinum*, with only *pe35/ppe68*, or with the *ppe68* operon of *M. tuberculosis* (*rv3872-rv3875*). Hemoglobin release was quantified by determining the OD_450_ of the medium after incubation.

To investigate the individual roles of *pe35/ppe68* and *esxB*/*esxA* in ESX-1 mediated secretion, we created an *M. marinum* knock-out strain that contained a deletion of the complete *pe35-ppe68-esxA-esxB* operon (Figure 1A). Again, no EspE was detected on the cell surface of the *ppe68* operon mutant, while the intracellular EspE levels were comparable to the WT (Figure 1C, lane 2, 9). Introduction of either *esxB/esxA* or *pe35/ppe68* on the integrative pMV vector failed to restore EspE and EsxA secretion (Figure 1C, lane 10-11). EspE and EsxA secretion was only restored when the entire *ppe68* operon was present on the introduced pMV or pSMT3 vector (Figure 1C, lane 12-13). Expression of the entire *ppe68* operon from the pSMT3 vector resulted in secretion levels similar to the WT. In addition, the *ppe68* operon of *M. tuberculosis* was also able to restore EspE and EsxA secretion by the *ppe68* operon mutant, showing that the function of the *ppe68* operon in ESX-1 secretion is conserved (Figure 1C, lane 14).

Next, we investigated the impact of the *ppe68* fs mutation on ESX-1 mediated membrane lysis, as an important readout for ESX-1 mediated virulence mechanisms (24). As expected, while WT *M. marinum* lysed sheep red bloods cells efficiently, a mutation in the ESX-1 core component EccC_b1_ showed no effect above background levels (Figure 1D). Also the *ppe68* fs mutant completely lost its hemolytic capabilities, which could be fully complemented by the introduction of the *ppe68* operon. Introduction of the *pe35/ppe68* pair on a pSMT3 plasmid and the *ppe68* operon of *M. tuberculosis* partially restored hemolysis (Figure 1C). In conclusion, *ppe68* is essential for the hemolytic activity of *M. marinum* and the secretion of EsxA and EspE, which role is conserved between *M. marinum* and *M. tuberculosis*.

### The EspG_1_-binding domain of PPE68 is important for its central role in ESX-1 secretion

Previously, we reported that the EspG_1_-binding domain of PPE68_1 (MMAR_0186) is essential for secretion via the ESX-1 system (43). Specifically, a single amino acid substitution at position 125, located within this domain, abrogated PPE68_1 secretion to the culture supernatant without affecting its production. Similarly, the conserved amino acid in an ESX-5 dependent PPE protein (L125) is essential for EspG_5_ binding and its secretion (11). To investigate the role of EspG_1_-binding to PPE68, an equivalent F125A amino acid substitution was introduced in the *ppe68* operon on the pSMT3 vector, with a Strep-tag to the C-terminus of PPE68 for detection purposes. Immunoblot analysis showed that PPE68.strep was properly produced and, as expected, ran slightly higher on an SDS-PAGE gel (Figure 2A, lane 4). Importantly, complementation of EsxA and EspE secretion by the *ppe68* operon mutant was not affected by the Strep-tag (Figure 2A, lane 12). While the F125A substitution did not affect the stability of this PPE68 variant (Figure 2A, lane 5), it could not restore WT levels of EsxA secretion and showed severely reduced EspE secretion in the *ppe68* operon mutant (Figure 2A, lane 13). This result indicates that recognition of PPE68 by EspG_1_ is indeed important for its role in ESX-1 secretion.

**Figure 2.**
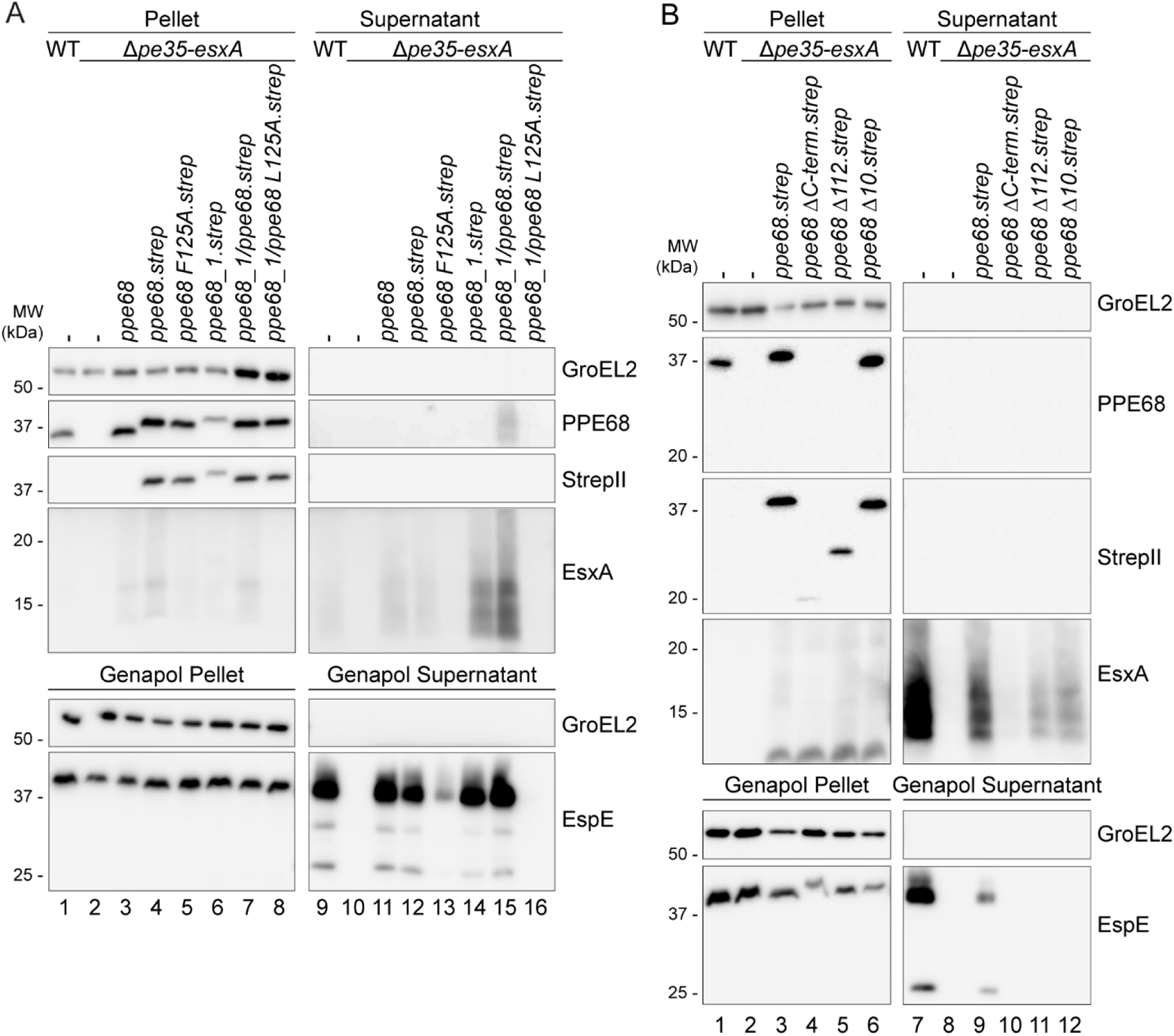
Conserved features of PPE68 are important for its central role in ESX-1 secretion. A) The role of the EspG_1_-binding domain of PPE68. SDS-PAGE and immunostaining of the cell pellet, culture supernatant, Genapol pellet and Genapol supernatant fractions of *M. marinum* WT, the *ppe68* operon mutant (Δ*pe35-esxA*) and the *ppe68* operon mutant complemented with either the *ppe68* operon, the *ppe68_1* operon, or with a hybrid operon containing *pe35_1*, the N-terminal domain of *ppe68_1*, the C-terminal domain of *ppe68*, *esxB* and *esxA*. Single amino acid substitutions were made in the helical tip of PPE68 and the PPE68_1/PPE68 hybrid to investigate the role of the EspG_1_-binding domain. B) The role of the C-terminal domain of PPE68. SDS-PAGE and immunostaining of the cell pellet, culture supernatant, Genapol pellet and Genapol supernatant fractions of *M. marinum* WT, the *ppe68* operon mutant (Δ*pe35-esxA*), and the *ppe68* operon mutant complemented with either the WT or *ppe68* operon variants, encoding PPE68 lacking either the entire (PPE68ΔC-term) or partial, i.e. the final 112 (PPE68Δ112) or 10 (PPE68Δ10) amino acids, C-terminal domain. In both A) and B), the operons were expressed from a multicopy pSMT3 vector under control by the *hsp60* promotor and with a Strep-tag on the C-terminus of PPE68 and its variants. Proteins were visualized with anti-EsxA, anti-EspE, anti-StrepII (PPE68) and anti-PPE68 antibodies (ESX-1 substrates). A processed band of 25 kDa was detected by the anti-EspE antibody in the Genapol supernatant fraction of the WT, which has been reported before (44). As a loading and lysis control, blots were incubated with antibodies directed against the cytosolic GroEL2 protein. In all blots, equivalent OD units were loaded: 0.2 OD for pellet and Genapol pellet and 0.4 OD for supernatant and Genapol supernatant fractions.

As we were unable to directly assess the secretion of PPE68, but previously could observe secretion of PPE68_1 (42, 43), we further investigated the paralogous *ppe68_1* operon, consisting of *pe35_1*, *ppe68_1*, *esxB_1* and *esxA_1* (Figure 1A). While *esxB_1* and *esxA_1* are almost identical to their paralogs in the *esx-1* gene cluster, *pe35_1* and *ppe68_1* show lower similarity, particularly for the C-terminal domains of the PPE proteins (Figure 1A and S2). Notably, deletion of this four-gene paralogous region did neither affect EspE secretion nor the hemolytic activity of *M. marinum*, showing that these endogenous genes are dispensable for these specific ESX-1 mediated processes (Figure S3). To assess the importance of the C-terminal domain of the PPE68 paralogs in ESX-1 secretion, we constructed a pSMT3 plasmid containing the *ppe68_1* operon (*mmar_0185-88*) and a *ppe68*/*ppe68_1* hybrid operon construct, containing *pe35_1*, the N-terminal domain of *ppe68_1* (PPE domain, residue 1-175, see Figure S2), the C-terminal domain of *ppe68* (encoding for residue 176-367), *esxA* and *esxB*. Again, a Strep-tag was introduced to the C-terminus of the PPE variants for detection purposes. Surprisingly, both the *ppe68_1* operon and the hybrid operon re-established EsxA and EspE secretion in the *ppe68* operon mutant (Figure 2A, lane 14-15), while the *ppe68_1* operon was also able to restore hemolytic activity in the *ppe68* fs mutant (Figure S3B). These results show that the paralogous, but divergent PPE68_1 is functionally similar to the PPE68 with regard to their role ESX-1 secretion. However, the fact that deleting the paralogous PPE68_1 region did not impact ESX-1 secretion indicates that this region is not expressed under the conditions used. Notably, while some low amounts of HA- or FLAG-tagged PPE68_1 could previously be detected in culture supernatants of *M. marinum* M (42, 43), PPE68_1 with a Strep-tag could not clearly be detected in this fraction (Figure 2A, lane 14). In contrast, the PPE68_1/PPE68 hybrid could readily be observed in the culture supernatant using the anti-PPE68 antibody (Figure 2A, lane 15). The anti-StrepII antibody failed to visualize this protein in this fraction, likely due to C-terminal processing of the PPE68_1/PPE68 hybrid upon secretion (see below). Most importantly, the introduction of an L125A mutation in the EspG_1_-binding domain of the PPE68_1/PPE68 hybrid completely blocked its ability to complement secretion of EspE and EsxA in the *ppe68* operon mutant and also abrogated the secretion of the PPE68_1/PPE68 hybrid itself (Figure 2A, lane 16). These combined results show that the amino acid in position 125, located within the helical tip of the EspG_1_-binding domain, is essential for the export of the PPE68/PPE68_1 hybrid, which in turn is necessary for the secretion EspE and EsxA.

### The C-terminal domain of PPE68 is necessary for ESX-1 mediated secretion

While the C-terminal domains of PPE68_1 and PPE68 are relatively dissimilar, they possess a well-conserved stretch of 15 amino acids consisting of an unusual high amount of negatively-charged residues at their C-termini. To investigate the role of the C-terminal domain of PPE68 in more detail, several C-terminal truncations were made, namely PPE68ΔC (residue 1-174), PPE68Δ112 (residue 1-256) and PPE68Δ10 (residue 1-357) (Figure S2), including a C-terminal Strep-tag and encoded by the pSMT3 vector together with *pe35*, *esxB* and *esxA*. Immunoblot analysis of the pellet fractions using the anti-Strep antibody showed that the two more severe C-terminal truncations of PPE68 were detected in lower amounts, and only the PPE68Δ10 variant was produced at similar levels as the full-length protein (Figure 2B, lane 4-6). The anti-PPE68 antibody could only recognize the PPE68Δ10 variant, indicating that the antibody primarily recognizes the C-terminal domain of this protein. PPE68Δ122 and PPE68Δ10 were able to restore EsxA secretion by the *ppe68* operon mutant (Figure 2B, lane 11-12), whereas with the PPE68ΔC variant EsxA secretion could hardly be observed, most-likely because this variant is not stably produced (Figure 2B, lane 10). In contrast, while the intracellular protein levels of EspE were similar between all strains, none of the PPE68 variants showed detectable secretion of EspE (Figure 2B, lane 4-6 and 10-12). These results suggest that the C-terminal domain of PPE68 is not required for EsxA secretion, but it is necessary for the secretion of EspE.

### PPE68 is secreted independently from EsxA/EsxB and processed on the cell surface

Although PPE68 contains all characteristics of an ESX substrate, we were not able to detect this protein in culture supernatants (Figure 1 and 2). However, when we obtained a concentrated cell surface protein extract from larger cultures that were additionally grown without Tween-80, which was shown to retain surface proteins (51), low levels of PPE68 were detected in the surface extract of WT *M. marinum* (Figure 3A, lane 8). Here, PPE68 seemed to be processed, as multiple bands were recognized by the anti-PPE68 antibody. Only full length PPE68 was observed by the anti-StrepII antibody, suggesting that the other bands detected by the anti-PPE68 antibody represent C-terminal truncates. C-terminal processing of the Strep-tag was also observed for the secreted fraction of the PPE68_1/PPE68 hybrid (Figure 2A). As we were now able to detect surface-localized PPE68, we analyzed whether the export of PPE68 is dependent on the co-expression of *esxB/esx*A. Importantly, a pSMT3 vector with only the *pe35/ppe68.strep* genes fully restored PPE68 secretion to the cell-surface by the two *ppe68* mutants (Figure 3A, lane 11, 14). As the *ppe68* operon mutant does not contain a genomic copy of *esxB*/*esxA*, these results suggest that PPE68 can be secreted independently of EsxA (Figure 3A, lane 14). Intriguingly, only full length PPE68.strep was detected on the cell surface of the complemented *ppe68* operon mutant, suggesting that surface-processing of PPE68 might be dependent on the co-secretion of EsxA.

**Figure 3.**
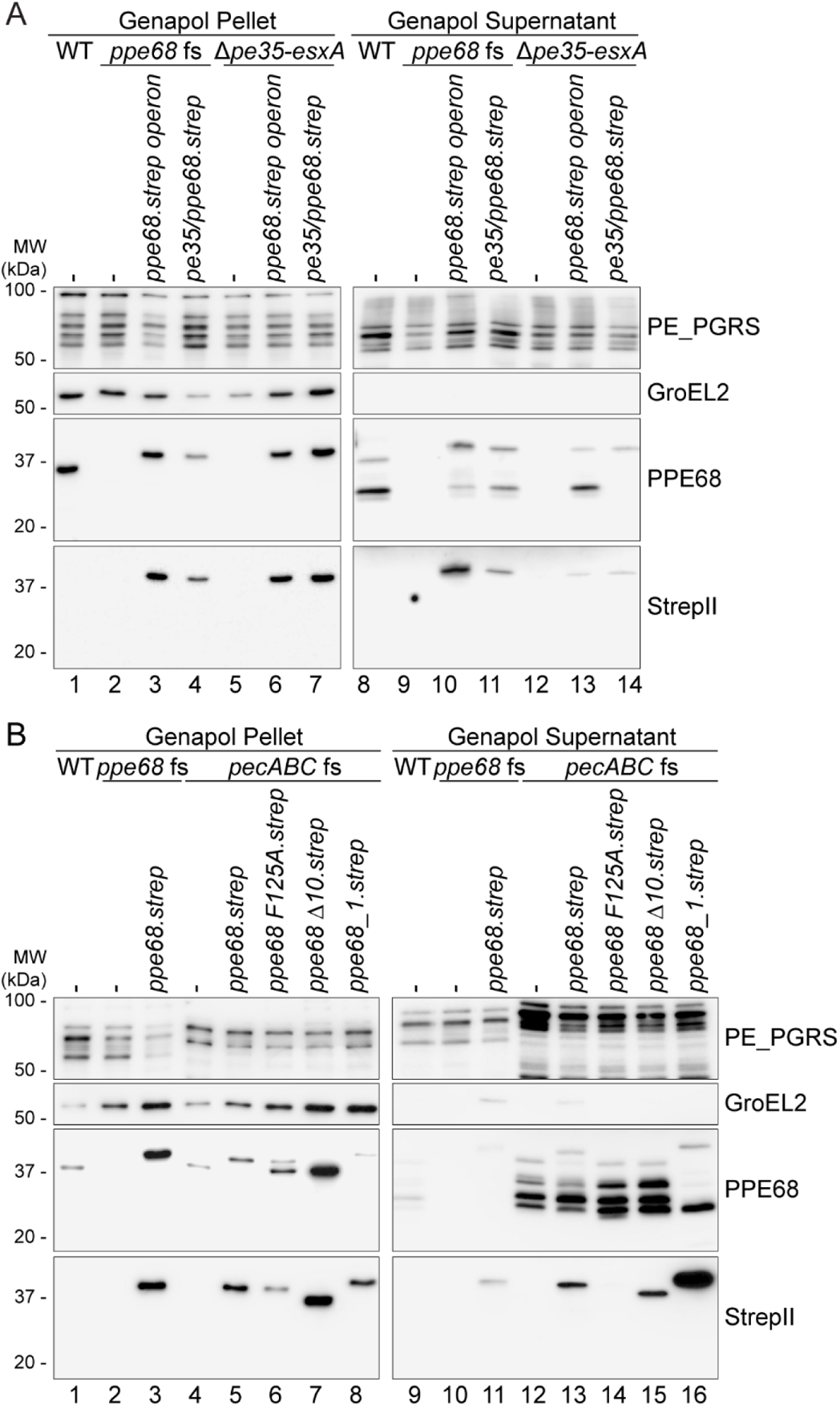
PPE68 is secreted independently of *esxB*/*esxA* and processed on the cell surface of *M. marinum*. A) SDS-PAGE and immunostaining on Genapol pellet and concentrated Genapol supernatant fractions of *M. marinum* WT, the *ppe68* fs mutant, the *ppe68* operon mutant and both mutants complemented with *pe35/ppe68* or the entire *ppe68* operon. B) Additional SDS-PAGE and immunostaining of Genapol pellet and concentrated Genapol supernatant fractions of the *pecABC* fs mutant overexpressing the WT *ppe68* operon (*ppe68.strep*), the operon producing either the PPE68F125A with an amino acid substitution in the helical tip, or PPE68Δ10 with a C-terminal truncation of the last 10 amino acids, or the *ppe68_1* operon (*ppe68_1.strep*). Operons were expressed from a multicopy pSMT3 vector controlled by the *hsp60* promotor and with a Strep-tag on the C-terminus of PPE68 and the variants. Proteins were visualized with anti-StrepII (PPE68) and anti-PPE68 antibodies (ESX-1 substrates). All *ppe68* fs and *pecABC* fs mutants contain an additional pMV TdTomato plasmid. As a loading and lysis control, blots were incubated with antibodies directed against the cytosolic GroEL2 protein and the surface-localized PE_PGRS proteins. In all blots, equivalent OD units were loaded: 0.2 OD for Genapol pellet and 2 OD Genapol supernatant fractions.

The various processed forms of PPE68 on the surface indicates that this protein is degraded upon secretion. Previously, the aspartic protease PecA was shown to cleave PE_PGRS proteins on the cell surface of *M. marinum* (18). We therefore hypothesized that this protein together with its paralogs PecB and PecC could be involved in cell surface processing of PPE68. Using a *pecA-pecB-pecC* triple fs mutant, created using CRISPR1-Cas9 gene editing (Meijers *et al*., manuscript in preparation), we could confirm that, while PPE68 levels were similar in the Genapol pellet fractions (Figure 3B, lane 1, 4), the amount of PPE68 protein that was detected in the upscaled cell surface extract was significantly higher in the *pecABC* fs mutant compared to that of the WT (Figure 3B, lane 9, 12). This shows that the amount of exported PPE68 is in fact higher than was previously assumed. As similar processing of surface-localized PPE68 was still observed in the *pecABC* fs mutant, another protease should be involved in this process. A prime candidate would be the serine protease MycP_1_, which is one of the conserved ESX-1 components. However, a point mutation in the active site of MycP_1_ (52) did not affect the surface processing of PPE68 (Figure S4).

Next, we exploited the increased amount of cell surface-localized PPE68 in the *pecABC* fs mutant to further investigate the requirement for PPE68 secretion. When we disrupted the interaction with EspG_1_ by the F125A mutation, PPE68.strep was not detected in the surface fraction (Figure 3B, lane 13), confirming the importance of the EspG_1_ interaction for secretion. Interestingly, deletion of the C-terminal 10 amino acids of PPE68, which did not majorly affect EsxA secretion but abrogated secretion of EspE, did not interfere with secretion of PPE68 itself (Figure 3B, lane 15). To summarize, PPE68 is exported to the cell surface of *M. marinum*, where it is subsequently degraded. Secretion of PPE68 depends on its interaction with EspG_1_, but is independent of co-expression of *esxB*/*esxA* and its final 10 amino acids.

### Intracellular PPE68 forms a complex with EspG_1_ and a PE protein

While low amounts of PPE68 could be extracted from the cell surface, a large portion of PPE68 remained cell-associated upon detergent treatment (Figure 3). In *M. tuberculosis*, PPE68 was shown to localize to the cell envelope fraction by subcellular fractionations (8, 53). However, when we fractionated *M. marinum* cells by high-pressure lysis followed by ultracentrifugation, we found the entire population of intracellular PPE68 to be present in the soluble fraction (Figure 4A). In an *eccC_a1_* deletion strain, production and solubility of PPE68 was similar as in the WT, while deleting the *espG_1_* chaperone gene drastically reduced the amount of PPE68. Exogenous expression of the entire *ppe68* operon in the *ppe68* operon mutant did not alter the solubility of PPE68 (Figure 4B). To test whether the observed difference in solubility between PPE68 homologs was species-specific or dependent on experimental procedures, we fractionated *M. marinum* expressing the *ppe68* operon of *M. tuberculosis* and the auxotrophic *M. tuberculosis* strain mc^2^6020 using the same procedure. In both cases, PPE68 of *M. tuberculosis* was again exclusively present in the soluble fraction, from which we conclude that the observed differences for PPE68 localization are a results of different protocols for cell lysis and subcellular fractionation.

**Figure 4.**
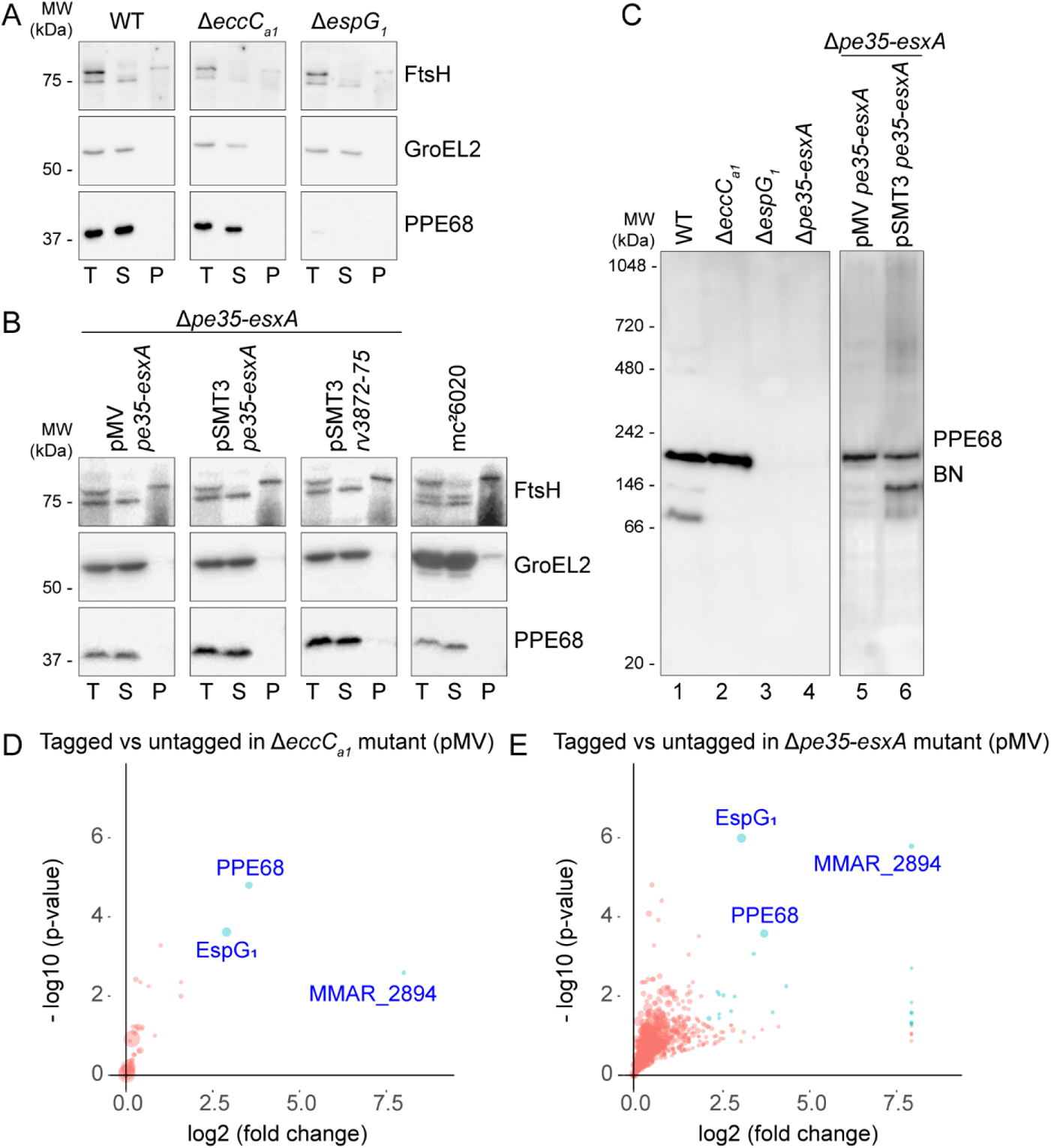
PPE68 of *M. marinum* and *M. tuberculosis* is soluble and forms complexes of different sizes. A,B) SDS-PAGE and immunostaining of subcellular fractionation of *M. marinum* WT, an *eccC_a1_* mutant and an *espG_1_* mutant (A) and the *ppe68* operon mutant complemented with the *ppe68* operon of *M. marinum* expressed from a pMV or pSMT3 vector, or the *ppe68* operon of *M. tuberculosis* H37Rv expressed from a pSMT3 vector, and the auxotrophic *M. tuberculosis strain mc^2^6020* (B). Proteins were visualized with anti-PPE68, while antibodies against GroEL2 (cytosolic protein) and FtsH (membrane protein) were used to assess the purity of the fractions. Notably, the FtsH antiserum cross-reacts with a cytosolic protein. In all blots, equivalent amounts corresponding to 0.2 OD units were loaded. T, total lysate, S, supernatant/soluble proteins, P, pellet/cell envelope proteins. C) Immunostaining of soluble fractions of the same strains as in A) and B) after blue native PAGE (BN-PAGE) gel electrophoresis. In all blots, equivalent amounts of 19 µg of total protein was loaded. D,E) PPE68 interacts with its PE partner, MMAR_2894, and the EspG_1_ chaperone in the cytosol. Quantitative proteomics analysis comparing the PPE68 affinity purification from the soluble fraction of the *eccC_a1_* mutant (D) and the *ppe68* operon mutant (E), complemented with the *ppe68* operon, producing PPE68 with or without a Strep-tag on its C-terminus from a pMV vector. Proteins that show log2 fold change > 2 and a −log10 p-value > 1.3 between tagged and untagged PPE68 are indicated as blue dots. Size of the dots are correlated to the number of spectral counts.

The current paradigm is that PPE proteins interact with a specific PE partner and a system-specific EspG chaperone in the cytosol prior to secretion (11, 12, 54). Analysis of PPE68 complex formation by blue native PAGE (BN-PAGE) revealed three major intracellular complexes ranging from ~70 to 180 kDa (Figure 4C, lane 1). Both the *ppe68* operon mutant and the Δ*espG_1_* mutant did not show any PPE68 complex formation (Figure 4C, lane 3-4). In the soluble fraction of the Δ*eccC_a1_* mutant, in which export of PPE68 is blocked, the largest (~180 kDa) and predominant complex was observed in similar amounts as in the WT fraction (Figure 4C, lane 2), suggesting that a large portion of PPE68 in WT bacteria is retained in the cytosol. The additional two complexes were absent in the Δ*eccC_a1_* mutant, indicating that these two assemblies are only formed in the presence of a functional ESX-1 membrane complex. Complementation the *ppe68* operon mutant with the *ppe68* operon of *M. marinum*, either using the pMV or pSMT3 plasmid, restored formation of the ~180 kDa PPE68 complex, while the two smaller complexes were only clearly observed when complemented with the pSMT3 vector (Figure 4C, lane 5-6). This difference could be due to higher expression levels of the operon genes of the latter vector, compared to that of the integrative pMV vector.

To analyze the composition of the observed complexes, a Strep-affinity purification was performed on the soluble fractions of the Δ*eccC_a1_* and *ppe68* operon mutant, producing Strep-tagged PPE68 from either the pMV or pSMT3 vector containing the full *ppe68* operon. Both mutants containing the *ppe68* operon without a Strep-tag sequence were included as negative controls. BN-PAGE analysis of the purified samples showed a similar pattern of PPE68 complexes as seen for endogenous PPE68 (Figure S5C). LC-MS/MS analysis showed that the PPE68.strep samples, purified from the Δ*eccC_a1_* mutant, contained, as expected, PPE68 and the chaperone EspG_1_. The third protein that was present in high amounts was an *M. marinum*-specific PE protein, named MMAR_2894 (Figure 4D and Table S1). Interestingly, this protein was already identified as an ESX-1 substrate (44). Notably, PE35 of *M. marinum* lacks, unlike its ortholog in *M. tuberculosis*, a general YxxxD/E secretion motif (Figure S6) (14), which might explain why PPE68 binds an alternative partner in *M. marinum*. As PE35_1 was also not detected, MMAR_2894 seems to be the only PE partner of PPE68 in *M. marinum*. EspG_1_ and MMAR_2894 were also the predominant co-purified proteins of PPE68 purified from the complemented *ppe68* operon mutant, confirming that intracellular PPE68 localizes to the cytosol also in the presence of a functional ESX-1 system (Figure 4E and Table S1). PPE68.strep produced from the pSMT3 vector in both the *ppe68* operon and Δ*eccC_a1_* mutant also showed the specific co-purification of EspG_1_ and MMAR_2894 (Figure S5D and E).

To confirm the presence of MMAR_2894 and EspG_1_ in the PPE68 complexes, we created an *mmar_2894* fs mutant and analyzed the PPE68F125A variant, respectively. The *mmar_2894* fs mutant, which contained a 13 bp deletion after 192 base pairs that resulted in an early stop codon after 68 amino acids (Figure S1B), showed an ESX-1 secretion defect and loss of hemolytic capabilities similar to the *ppe68* fs mutant (Figure S7A-C). While PPE68 was detectable and soluble in the *mmar_2894* fs mutant, BN-PAGE analysis showed that the major ~180 kDa complex was absent in this mutant. The smaller two complexes were still present, suggesting only the predominant complex contained MMAR_2894 (Figure 5B, lane 3). With the PPE68F125A variant, no complexes could be observed by BN-PAGE analysis (Figure 5B, lane 5), while it was detectable and in the soluble fraction by SDS-PAGE analysis, suggesting that all three complexes contain EspG_1_.

**Figure 5.**
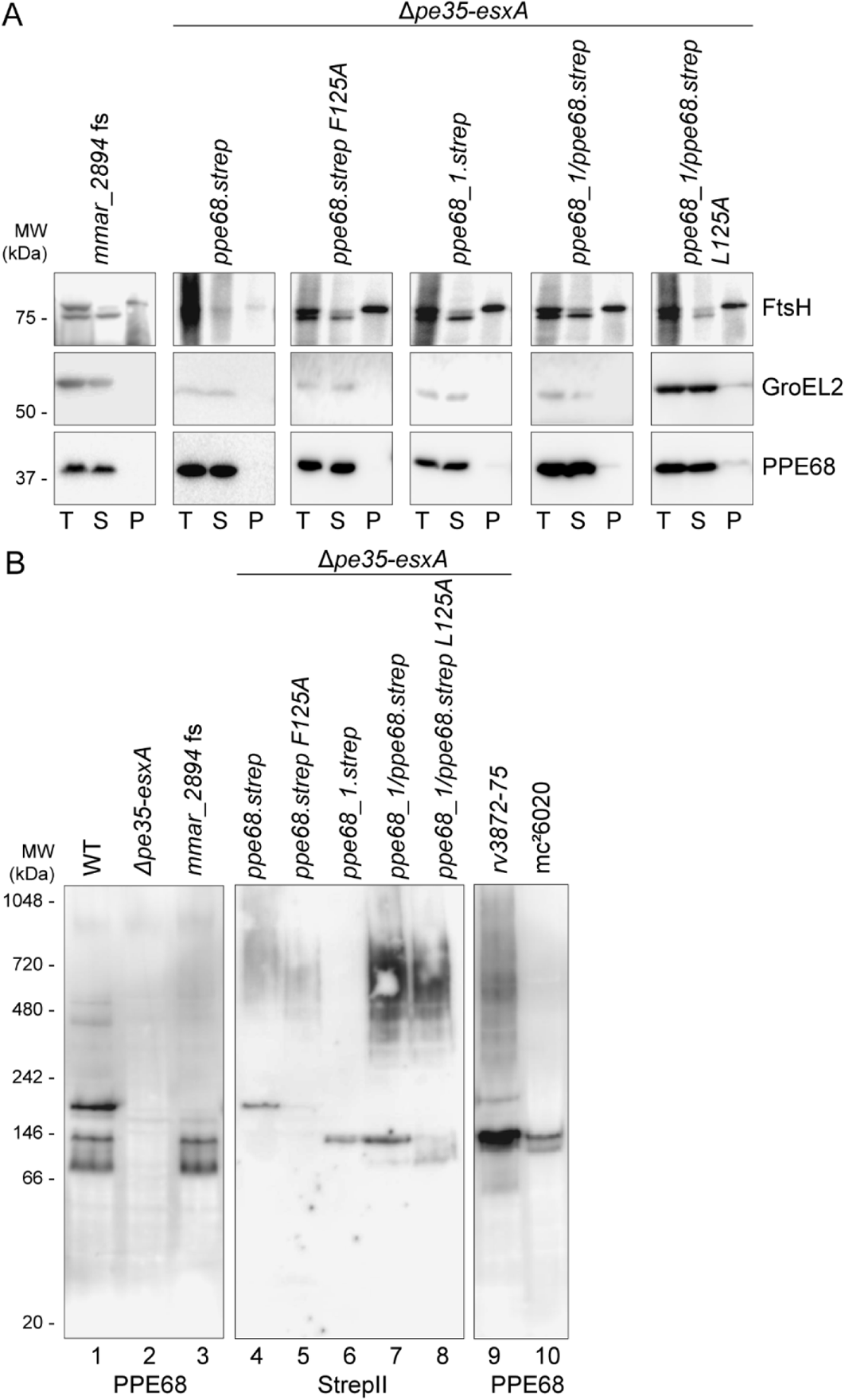
Verification of complex formation of PPE68 with MMAR_2894 and EspG_1_. A) SDS-PAGE and immunostaining of subcellular fractions of the *mmar_2894* fs mutant and the *ppe68* operon mutant complemented with the WT *ppe68* operon, the *ppe68_1* operon, a hybrid operon containing *pe35_1*, the N-terminal domain of *ppe68_1*, the C-terminal domain of *ppe68* and *esxB* and *esxA* or the *ppe68* or hybrid operon expressing a PPE68 variant with a single amino acid substitution at position 125. Genes were expressed from a multicopy pSMT3 vector under the *hsp60* promotor with a Strep-tag on the C-terminus of PPE68 or its variants. Proteins were visualized with anti-PPE68 antibodies, while antibodies against GroEL2 (cytosolic protein) and FtsH (membrane protein) were used to assess the purity of the fractions. Notably, the FtsH antiserum cross-reacts with a cytosolic protein. In all blots, equivalent amounts corresponding to 0.2 OD units were loaded. T, total lysate, S, supernatant/soluble proteins, P, pellet/cell envelope proteins. B) BN-PAGE and immunostaining of soluble fractions of the same strains as in A, and also the *ppe68* operon mutant expressing the *ppe68* operon of *M. tuberculosis* from a pSMT3 vector and the auxotrophic *M. tuberculosis* strain mc^2^6020. Proteins were visualized with anti-PPE68 and anti-StrepII antibodies. In all blots, equivalent amounts of 19 µg of total protein was loaded.

Finally, we also analyzed complex formation of PPE68 from *M. tuberculosis* using both heterologous expression of the *ppe68* operon in *M. marinum* and the auxotrophic *M. tuberculosis* strain mc^2^6020. The observed PPE68 complexes in both strains have the same size as PPE68_1.strep and the PPE68_1/PPE68.strep hybrid. The smaller size of these complexes compared to the ~180 kDa PPE68_mmar_ complex could be due to the sizes of their most likely PE partners, i.e. PE35_1 and PE35_mtub_, respectively, which are smaller (9 kDa) than MMAR_2894 (22 kDa). Structural analysis suggests that PE-PPE-EspG complexes are formed by a single copy of each protein (11, 12), resulting in a molecular weight for MMAR_2894-PPE68-EspG_1_ of *M. marinum*, and both PE35_1-PPE68_1-EspG_1_ of *M. marinum* and PE35-PPE68-EspG_1_ of *M. tuberculosis* of 89 kDa and 74 kDa, respectively. The fact that the observed complexes have twice the molecular weight, i.e. ~180 kDa and ~150 kDa, respectively, indicate that the PE-PPE-EspG complexes form a dimer of trimers in the cytosol of mycobacteria. In summary, our fractionation and pulldown results suggest that cell-associated PPE68 of *M. marinum* and *M. tuberculosis* is mainly located in the cytosol, where it is in a complex together with EspG_1_ and its PE partner.

### The role of PPE68 in ESX-1 secretion: a model

Based on our results we propose a working model for PPE68 in the secretion of EsxA and EspE via the ESX-1 secretion system in *M. marinum* (Figure 6). In the cytosol PPE68 forms a stable complex together with its PE partner, *i.e.* MMAR_2894 in *M. marinum*, and chaperone EspG_1_. Upon targeting to the ESX-1 membrane complex, the PE/PPE pair binds to the central ATPase EccC_ab1_, possibly to the linker 2 domain between nucleotide-binding domain (NBD) 1 and 2, a previously proposed recognition site for PE/PPE substrates (55). Based on our observation that PPE68 does not require EsxA/EsxB for its secretion, this binding event is enough to trigger activation of the translocation channel. As a second step, EsxA/EsxB binds, based on *in vitro* binding studies and structural analysis, to NBD3 of EccC_ab1_ (23, 56–58). The Esx pair probably also interacts with the PE/PPE pair at the membrane complex, as PE35_1/PPE68_1 has been shown to determine the system-specificity of EsxB_1/EsxA_1 (42). As the third step, EspE is targeted with its predicted partner EspF and chaperone EspH (44) to the same ESX-1 complex. Secretion of the PE/PPE and Esx pairs probably does not depend on EspE, as mutations in EspH only affects secretion of EspE and EspF (44). Subsequently, the three substrate pairs are transported together through the channel at the cost of ATP hydrolysis by the EccC ATPases. EspE interacts with the C-terminus of PPE68 during the translocation process, as these residues are important for the secretion of specifically this Esp substrate. The mechanism of translocation across the mycobacterial outer membrane remains unknown. While it is tempting to speculate that PPE68 plays a role in the process, the protein does not form a stable channel in this specific membrane. Instead, the main portion of PPE68 exists as a cytosolic pool, where it serves as a reservoir to facilitate secretion of the other ESX-1 substrates. Once PPE68 is exported to the cell surface, it is processed and/or degraded by PecABC and an unspecified protease.

**Figure 6.**
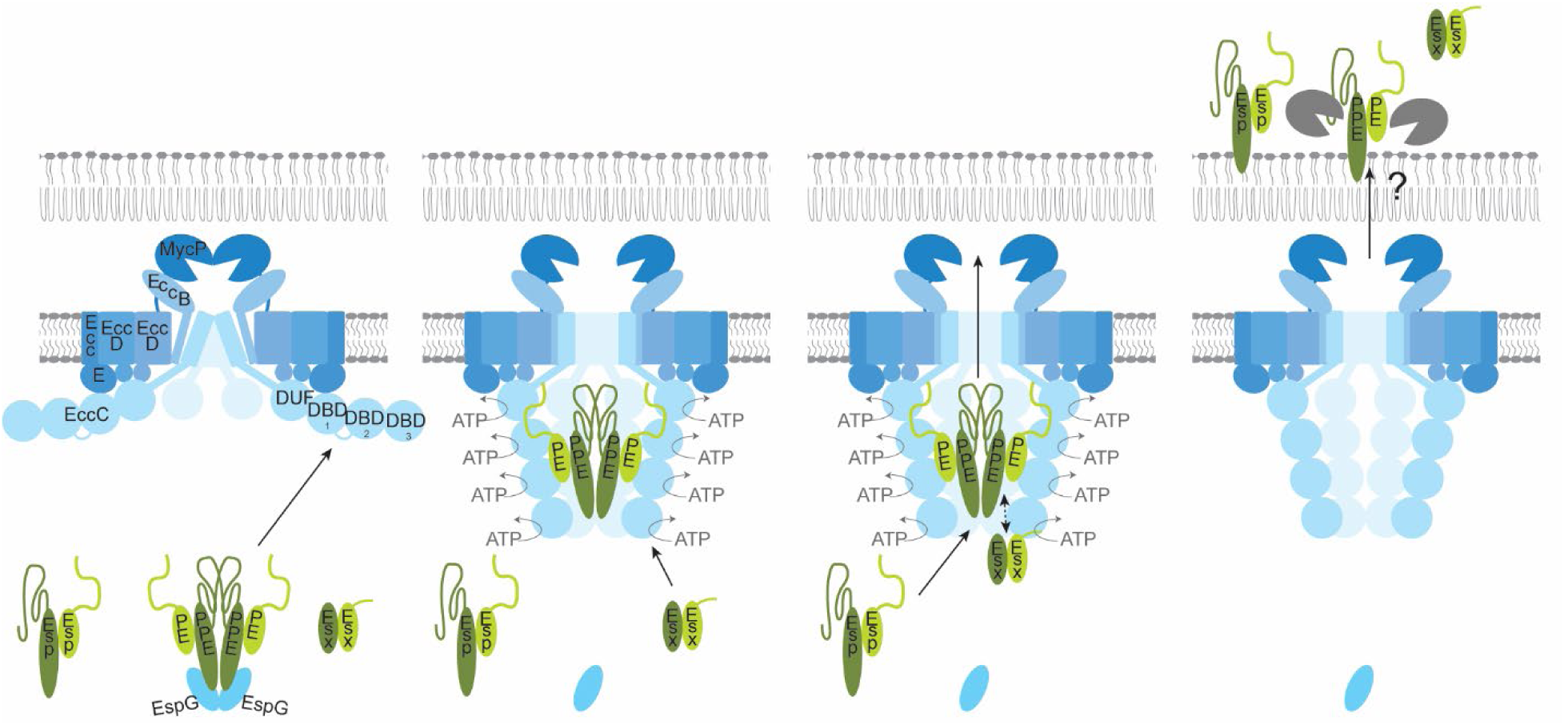
Working model for the role of PPE68 in ESX-1 mediated secretion. See text for an explanation.

## Discussion

The ESX-1 substrate EsxA is an abundantly secreted and highly immunogenic protein of *M. tuberculosis* and *M. marinum* (59–61) and has therefore been a major focus to get mechanistic insight into the essential role of ESX-1 in phagosomal membrane rupture by these pathogens (2–4, 22–24). However, the investigation of EsxA, or other ESX-1 substrates, remains difficult due to the co-dependent secretion of these substrates (32, 35–38). Here, we report that ESX-1 substrate PPE68, which is encoded within the same operon as EsxA, plays a central role in the secretion of EsxA and EspE in *M. marinum* and provide mechanistic insight into the observed co-dependent secretion of these substrates.

First, we showed that PPE68 is exported in an ESX-1 dependent manner to the cell surface of *M. marinum*. While previous mass spectrometry analysis could also detect this protein in culture supernatants of *M. marinum* (34, 44), we could not detect PPE68 in similar fractions by immunoblot analysis. Intriguingly, this protein is degraded upon secretion to the cell surface, as a deletion of three Pec proteases (PecABC) increases the amount of PPE68 on the cell surface. This provides an explanation for the low amount of exported PPE68 we detected. The observation that the *pecABC* deletions did not affect the processing pattern of PPE68 protein suggests that an additional protease is involved in its processing/degradation.

A frameshift mutation in *ppe68* abolishes secretion of EsxA and EspE, and hemolytic activity by *M. marinum*, which could be fully complemented by introduction of the *pe35*-*ppe68*-*esxB*-*esxA* operon. The role of PPE68 in the secretion process depends on the export of the PPE protein, as a point mutation in its EspG_1_-binding domain that blocks PPE export, abolishes secretion of the other ESX-1 substrates. Importantly, PPE68 export is independent of *esxA*/*esxB* expression, showing there is a hierarchy in co-dependent secretion. While the Esx pair has been considered to be important for secretion of all other ESX substrates, this is, to our knowledge, the first demonstration of an ESX-1 substrate that is fully independent of *esxB*/*esxA* for its export. Notably, we showed that PPE68 of *M. tuberculosis* can take over the central role of endogenous PPE68 of *M. marinum* in ESX-1 mediated secretion and hemolysis. This contrasts previous studies, showing that interruption of the *ppe68* gene did not affect secretion of EsxA in *M. tuberculosis* H37Rv (53, 62), although similar mutations did affect growth of these strains in a mouse infection model (62, 63). While the reason for the observed differences in requirements for EsxA secretion remains unclear, these might be caused by a variation in *in vitro* growth conditions, similarly as previously observed for the importance of the conserved cytosolic component EccA_1_ in secretion (44). Notably, a recent proteo-genetic study also revealed the importance of PPE68 and MMAR_2894 for ESX-1 mediated secretion in *M. marinum* (64), fully supporting our results.

A co-dependent relationship between substrates of the same ESX system could be explained by a more integral role of specific substrates in the translocation process itself. Interestingly, PPE68 of *M. tuberculosis* has previously been shown to localize to cell envelope fractions (8, 53), which could suggest a role in substrate translocation across the second mycobacterial membrane. However, we observed that cell-associated PPE68 is soluble, both in *M. marinum* and *M. tuberculosis,* which is not expected for an outer membrane channel. These contrasting observations are probably caused by experimental differences. Although a function for PPE68 in outer membrane translocation is still possible, the fact that we find this protein predominantly in the cytoplasmic compartment points towards a role in substrate recognition and/or activation by the ESX-1 inner membrane complex. Our previous observation that PPE68_1 determines through which ESX system EsxA_1 is exported by *M. marinum* is in agreement with such a function (42). In addition, a domain in the EccC ATPase component, *i.e.* linker2 between NBD_1_ and NBD_2_, has previously been shown to be involved in species-specific secretion of PE_PGRS proteins via the ESX-5 system (55). We therefore propose that binding of the PE/PPE pair to linker2 on EccC_ab1_ could trigger the activation of the ATPases and the opening of the ESX inner membrane complex.

Interestingly, we observed that the final 112 amino acids of PPE68 are dispensable for EsxA secretion. This suggests that the conserved N-terminal PPE domain of this protein, probably together with the PE domain of its partner, is involved in the secretion of other ESX-1 substrates. This involvement could be indirect, by activation of the ESX membrane channel, but could also be mediated by a direct interaction of the two substrate pairs. In contrast, the C-terminus of PPE68 seems to be required for the secretion of EspE, i.e. deletion of only the C-terminal ten amino acids already blocked EspE secretion. This charged C-terminal tail is conserved in the PPE68 homolog of *M. tuberculosis* and the PPE68_1 paralog in *M. marinum,* possibly explaining why these homologs are able to complement EspE secretion in the *M. marinum ppe68* mutants. We propose that EspE interacts with the C-terminal domain of PPE68 during the secretion process.

The major portion of cytosolic PPE68 is in a ~180 kDa complex with the EspG_1_ chaperone and an unexpected PE protein, namely MMAR_2894. MMAR_2894 has previously been identified as an ESX-1 substrate with a role in hemolysis (44, 65). Mutating *mmar_2894* leads to the absence of the major cytosolic PPE68 complex and a secretion and hemolytic defect identical to the *ppe68* fs mutant, which confirms that MMAR_2894 is the PE partner of PPE68 in *M. marinum*. This also means that PPE68 does not pair with the adjacently encoded PE35, which explains why PE35 lacks a T7SS signal (YxxxD/E). Apparently, during evolution PE35 was replaced by MMAR_2894 and lost its function, or *vice versa*. Analysis of multiple available *M. marinum* genomes (66) revealed a widespread lack of the T7SS signal in PE35, which extends to its close relatives *M. liflandii* and *M. ulcerans,* suggesting that this variation arose in a common ancestor. Seemingly, new associations between PE and PPE proteins can be formed, which also cautions the interpretation of adjacent PE and PPE genes as necessary secretion partners. Based on available structural data on PE:PPE:EspG complexes and observed molecular weight of the major cytosolic complex, we postulate that the MMAR_2894:PPE68:EspG_1_ complex forms a dimer of trimers. Dimers of EspG_3_ from *M. tuberculosis* and *M. smegmatis* have been crystalized, suggesting that dimerization could occur via the chaperone (12, 67).

Overall, our study describes a central role for PPE68 in ESX-1 mediated secretion and function. We hypothesize that PPE68 is in complex together with its PE partner and EspG chaperone in the cytosol as a reservoir to aid in the secretion of other substrates across the inner and possibly also the outer membrane. Further studies should focus on the exact mechanism of co-dependence of substrates for secretion, potential interactions between secreted co-dependent heterodimers and the role of the EccC ATPase in this process. The recent publications of the high-resolution structure of various ESX systems (68–71) will stimulate the field towards a mechanistic understanding of T7SS.

## Material and Methods

### Bacterial strains and culture conditions

All strains included in this study are listed in Table S2. Mycobacterial strains were routinely grown on Middlebrook 7H10 agar with 10% OADC (Difco) or in Middlebrook 7H9 liquid medium with 10% ADC (Difco) and 0.05% Tween-80. Additional supplement was necessary for culturing of *M. tuberculosis* mc^2^6020 (Δ*lysA* Δ*panCD*), containing 100 ng/ml L-lysine (Sigma) and 25 ng/ml D-panthotenic acid (Sigma). Appropriate antibiotics were added when necessary, i.e. 50 μg/ml kanamycin (Sigma), 50 μg/ml hygromycin (Roche) or 30 μg/ml streptomycin (Sigma). *M. marinum* and *M. tuberculosis* cultures and plates were incubated at 30 °C and 37 °C, respectively. *E. coli* DH5α was used for cloning and was grown on LB agar plates at 37 °C. Antibiotics were added at the same concentrations as described above.

### Plasmid construction

Genes of interest were amplified from *M. marinum* M or *M. tuberculosis* H37Rv genomic DNA by polymerase chain reaction (PCR) using primers that were synthesized by Sigma (Table S3). Hybrid combinations of genes or introduction of affinity tags were achieved with the use of nested primers. The Phusion polymerase (New England Biolabs, NEB) was used throughout all cloning procedures. Generated constructs were digested with XmnI and HindIII or NheI and BamHI for ligation into pMV361 and pSMT3 vectors (72, 73), respectively. Restrictions and ligation enzymes were provided by New England Biolabs (NEB). All plasmids were verified by Sanger sequencing (Macrogen) and are listed in Table S2.

### Generation of knockout strains and frameshift mutants

Generation of the *mmar_5447-50* and *mmar_0185-88* knockout strains were produced in *M. marinum* M by allelic exchange using a specialized transducing mycobacteriophage as previously described (74). PCR was used to amplify the flanking regions of the genomic regions. Deletions were confirmed by PCR, after which the selection markers were removed with a temperature sensitive phage encoding the γδ-resolvase (TnpR) (a kind gift from Apoorva Bhatt, University of Birmingham, UK). The frameshift (fs) mutants were obtained by genome editing using *Streptococcus thermophilus* CRISPR1-Cas9 (46). Single guide RNAs were produced by the annealing of oligonucleotides containing BsmBI overhangs (75). sgRNA were ligated into BsmBI digested pCRISPRx-Sth1Cas9-L5. The plasmids were sequenced and subsequently electroporated into competent *M. marinum* M cells. Bacteria were plated on 7H10 plates containing kanamycin and 100 ng/ml anhydrotetracycline (ATc, IBA Life sciences) for the induction of the *S. thermophilus* CRISPR1-Cas9 system. Single colonies were picked and screened for mutations in the gene of interest by PCR and sequencing. After verification, the CRISPR1-Cas9 integrative plasmid was removed by exchanging it with pTdTomato-L5 that integrates at the same site.

### RNA isolation and quantitative RT-PCR

RNA was isolated from bacterial cultures grown to an OD_600_ of 1. Bacteria were lysed by bead-beating in the presence of buffer RA-1 (NucleoSpin RNA isolation kit, Mackery-Nagel) and β-mercaptoethanol (Sigma). RNA isolation and purification was continued according to the protocol supplied by the manufacturer (Nucleospin RNA isolation kit, Mackery-Nagel). Samples were eluted in water with Ribolock (Thermoscientific) and an additional DNase treatment was performed with DNaseI (Thermoscientific) according to the protocol supplied by the manufacturer. cDNA was synthesized from 400 ng RNA with the RevertAid First Strand cDNA Synthesis Kit (Thermoscientific). Prior to the PCR reactions the cDNA was diluted 1:10 in RNase free water. Reaction were set up using iTaq Universal SYBR® Gren Supermix (Biorad). Quantitative RT-PCR was performed with the QuantStudio™ 3 Real-Time PCR System (ThermoFisher). At the end of amplification, PCR product specificity was verified by melt curve analysis and agarose gel electrophoresis. The threshold cycle (Ct) values were normalized with the Ct of SigA.

### Protein secretion

*M. marinum* cultures were grown until mid-logarithmic phase (OD_600_ of 1–1.4), before they were washed with 7H9 liquid medium containing 0.2% dextrose and 0.05% Tween-80 to clear the cultures of Bovine Serum Albumin (BSA). Cultures were set at an OD_600_ 0.4-0.5 and grown for 16 h. Cells were pelleted (10 min at 3,000xg), washed and resuspended in Phosphate-buffered saline (PBS) before lysing by bead-beating and 1/4^th^ volume of 5x SDS loading buffer (312.5 mM Tris-HCl pH 6.8, 10% SDS, 30% glycerol, 500mM DTT and 0.2% bromophenol blue) was added, yielding the pellet fraction. Culture medium was passed through an 0.22-µm filter and proteins were precipitated with Tricholoacetic acid (TCA). TCA precipitated pellets were washed with acetone and resuspended in 1x SDS loading buffer, yielding the supernatant fraction. Alternatively, a fraction of pelleted bacteria was incubated with 0.5% Genapol X-080 in PBS for 30 min with rotation at room temperature to extract cell surface associated proteins. Genapol-treated cells were pelleted, washed and resuspended in PBS before lysis by bead-beating and addition of 1/4^th^ volume of 5x SDS buffer, yielding the Genapol pellet fraction. 1/4^th^ volume of 5x SDS buffer was also added to Genapol supernatants, yielding the Genapol supernatant fraction. All fractions were boiled for 10 minutes at 95°C before loading on SDS-PAGE gels.

### Fractionation and Strep-tagged protein purification

Mycobacterial cultures were grown until an OD_600_ of 1. Cells were harvested and washed with PBS before resuspension in PBS with 10% glycerol to a concentration of 50-100 OD/ml. Cells were lysed by passage through a One-Shot cell disrupter (Constant Systems) at 0.83 kbar. Unbroken cells were pelleted by centrifugation at 15,000xg for 10 min at 4 °C. The obtained supernatant was subjected to ultracentrifugation at 100,000xg for 1hr at 4 °C, yielding the cell envelope pellet and soluble fractions. Samples were boiled in SDS buffer for 10 minutes at 95°C before loading on SDS-PAGE, as explained above. Alternatively, soluble fractions were then separated under native conditions on a 4–16% NativePage Novex BisTris Protein Gel (Life Technologies) and transferred to a polyvinylidene difluoride (PVDF) membrane before immunostaining.

For the purification of intracellular Strep-tagged proteins, cells were harvested and lysed in buffer B (PBS with 10% glycerol pH 8). Soluble fractions were prepared and incubated with StrepTactin resin (IBA Lifesciences) for 30 min with rotation at 4 °C. Beads were then washed with buffer B and proteins were eluted using buffer B supplemented with 10 mM desthiobiotin (IBA Lifesciences). Samples were taken at each step of the purification and loaded on SDS-PAGE gels to check the efficiency of affinity purification. Elution fractions were also separated under native conditions to ascertain the stability of the complexes after affinity purification.

### Protein detection and sera

SDS-PAGE and NativePage gels were stained with Brilliant Blue G-250 or R-250 (CBB; Bio-Rad), respectively. Alternatively, proteins were transferred to nitrocellulose (SDS-PAGE) or PVDF (BN-PAGE) membranes. Proteins were visualized using standard immunodetection techniques. Primary antibodies used in this study were anti-GroEL2 (CS44, Colorado state university), anti-PE_PGRS antibody (7C4.1F7) (76), anti-EsxA (Hyb76-8) (77), polyclonal anti-FtsH (78), polyclonal anti-PPE68 (8), polyclonal anti-EspE (50, 51) and polyclonal anti-Strep (Novusbio).

### Mass spectrometry

To identify the interaction partners of PPE68, elution fractions of the Strep-tagged protein purifications were analyzed by LC-MS/MS, essentially as described before (79). In short, SDS loading buffer was added to elution samples of Strep-tagged protein purification and boiled for 10 minutes at 95°C before loading on SDS-PAGE. SDS-PAGE gels were stained by CBB and total protein lanes were excised, washed and processed for in-gel digestion. Peptides were eluted from the gel pieces and analyzed by the Ultimate 3000 nanoLC-MS/MS system (Dionex LC-Packings, Amsterdam, The Netherlands). Proteins were identified from the resulting LC-MS/MS spectra by searching against the Uniprot *M. marinum* complete proteome (ATCC BAA-535M).

### Hemolysis assay

*M. marinum* strains were grown in 7H9 medium supplemented with ADC and 0.05% Tween 80 until mid-logarithmic phase. All strains were washed with PBS and diluted in Dulbecco’s Modified Eagle Medium (DMEM) without phenol red (Gibco, Life technologies). Number of bacteria were quantified by absorbance measurement at OD_600_ with an estimation of 2.5×10^8^ bacteria in 1 ml of 1.0 OD_600_. Defibrinated sheep erythrocytes (Oxoid-Thermo Fisher, the Netherlands) were washed and diluted in DMEM. A total of 4.15×10^7^ erythrocytes and 1.04×10^8^ bacteria (1:2.5 ratio) were added for one reaction of 100 μL in a round-bottom 96 well-plate, gently centrifuged for 5 minutes and incubated at 32°C with 5% CO_2_. After 3 hours of incubation, cells were resuspended and centrifuged. Supernatants were transferred to a flat-bottom 96-wells plate and measured at an absorbance of 405 nm to quantify hemoglobin release.

## Acknowledgements

We would like to thank Roy Ummels, Dagmar van Mourik, Daphne Fluitman and Jobana Ananthasabesan for their contributions to plasmid design and the generation of mutants. We are grateful to Eric Brown (Genentech) and Roland Brosch (Institut Pasteur) for offering anti-EspE and anti-PPE68 antibodies, respectively. This work was funded by the Netherlands Organization of Scientific Research through an ALW Open grant (ALWOP.319; to MPMD) and a VIDI grant (864.12.006; ENGH). The funders had no role in study design, data collection and analysis, decision to publish, or preparation of the manuscript

## Supplemental Information

**Figure S1.**
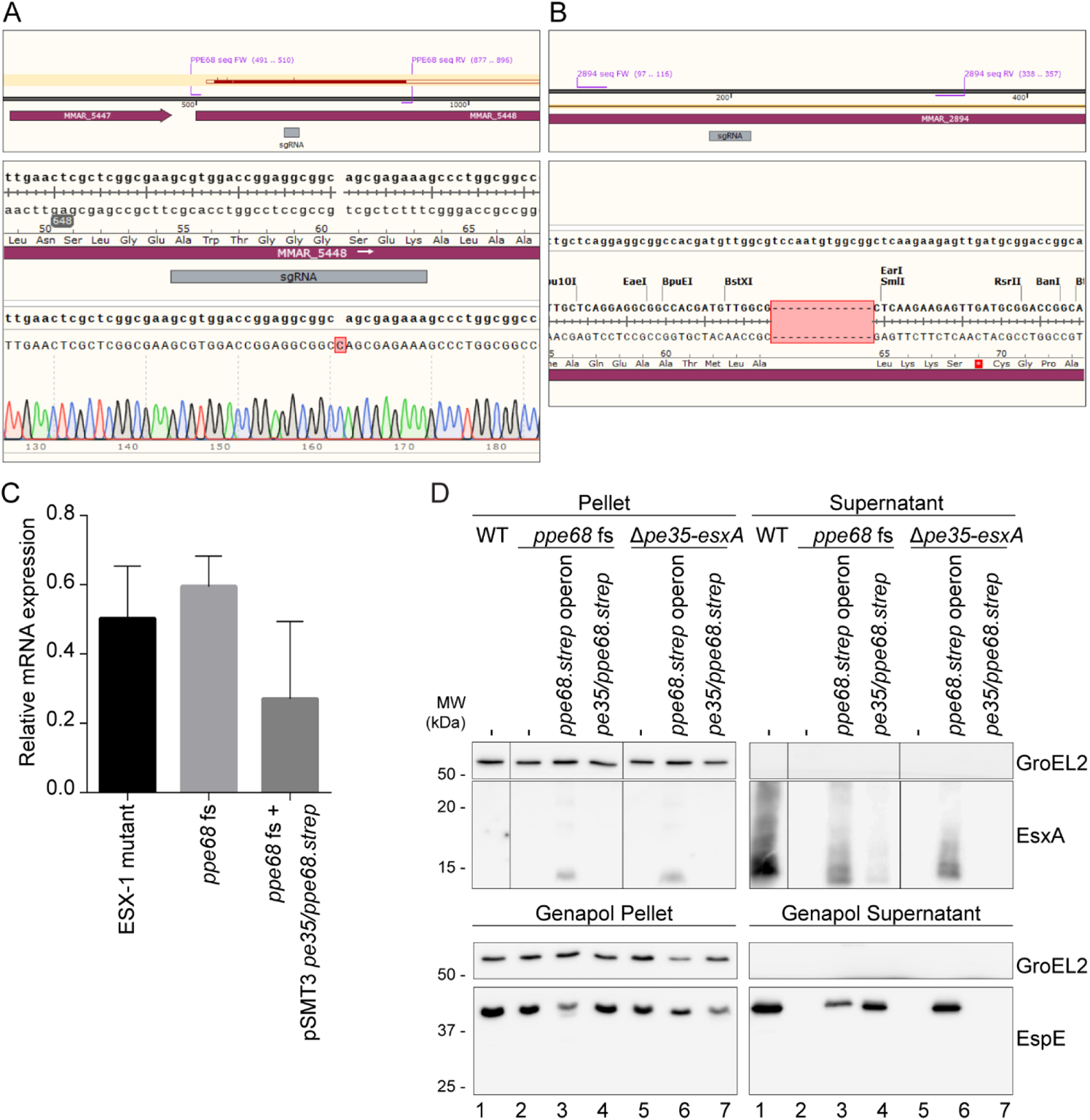
Generation and confirmation of *M. marinum* frameshift mutants, including phenotypic analysis of the *ppe68* fs mutant. A,B) Each panel shows the sgRNA binding region within the gene, *i.e.* in *ppe68* (A) and *mmar_2894* (B), with a panel underneath showing the sequencing results that were obtained after PCR. C) Relative *esxA* mRNA levels of an *eccC_b1_* mutant (M^VU^),the *ppe68* fs mutant and the *ppe68* fs mutant complemented with *pe35*/*ppe68* expressed from the pMST3 vector, compared to the WT. Cycle threshold (Ct) values were normalized for Ct values of the household gene *sigA* and compared to the Ct values. D) SDS-PAGE and immunostaining of the cell pellet, culture supernatant, Genapol pellet and Genapol supernatant fractions of *M. marinum* WT, the *ppe68* fs mutant with and without various a complementation constructs and the *ppe68* operon mutant (Δ*pe35-esxA*) with and without various complementation constructs. Genes were expressed from a multicopy pSMT3 vector under control of the *hsp60* promotor with a Strep-tag on the C-terminus of PPE68. The WT, the *ppe68* fs mutant and the *ppe68* fs mutant containing the pSMT3 vectors also contained a pMV TdTomato plasmid. Proteins were visualized with anti-EsxA and anti-EspE (ESX-1 substrates) antibodies. As a loading and lysis control, blots were incubated with antibodies directed against the cytosolic GroEL2 protein. In all blots, equivalent OD units were loaded: 0.2 OD for pellet and Genapol pellet fractions, 1.2 OD for supernatant fractions and 2 OD for Genapol supernatant fractions.

**Figure S2.**
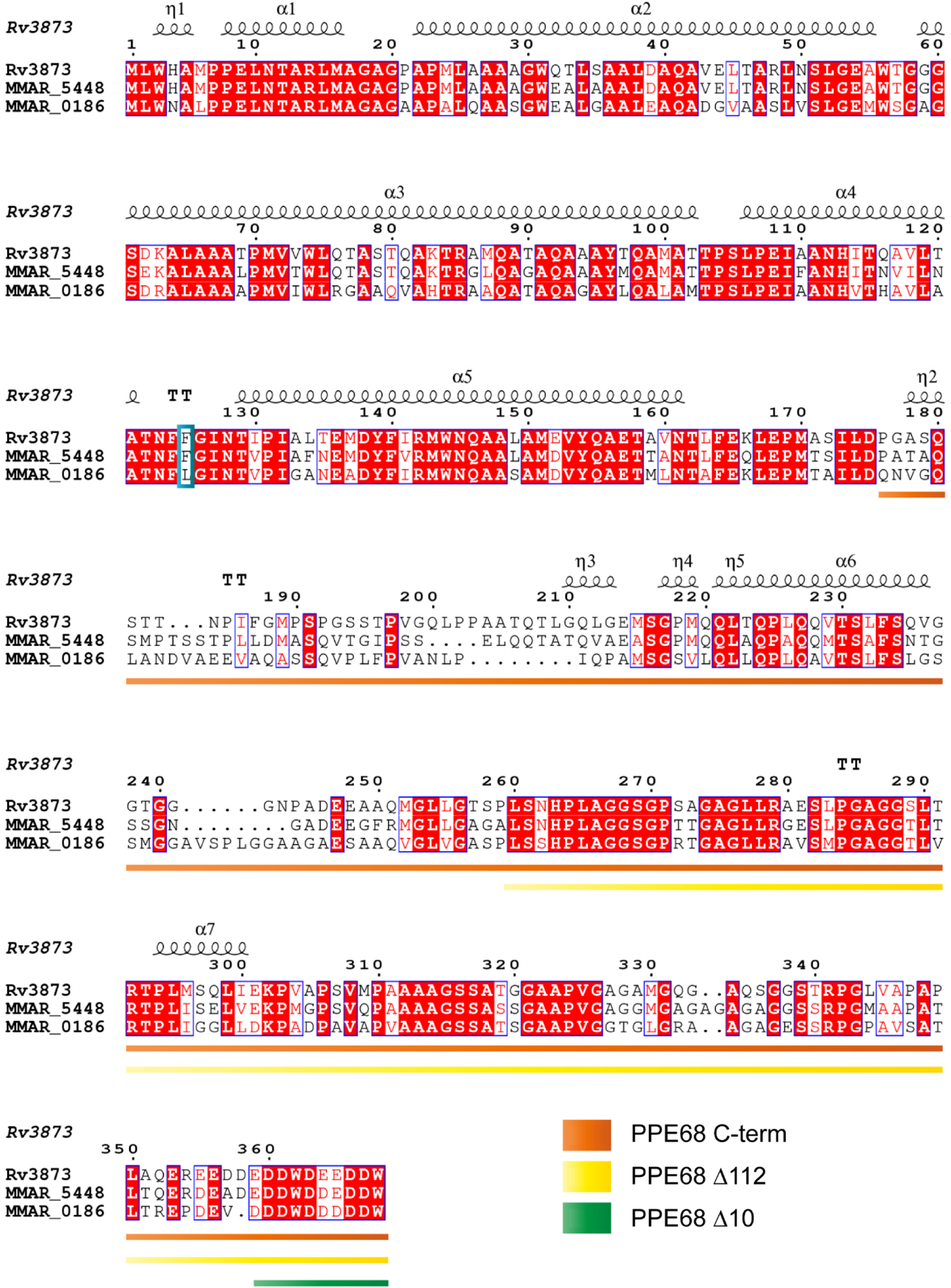
Protein sequence alignment of PPE68 homologs of *M. marinum* and *M. tuberculosis* including the secondary structure prediction of *M. tuberculosis* PPE68. The secondary structure prediction was produced by Alphafold. The blue square indicates the residue at position 125 involved in EspG_1_-binding. Underlined sequences indicate the C-terminal truncations according to the color key. Rv3873 = PPE68 of *M. tuberculosis*, MMAR_5448 = PPE68 of *M. marinum* and MMAR_0186 = PPE68_1 of *M. marinum*.

**Figure S3.**
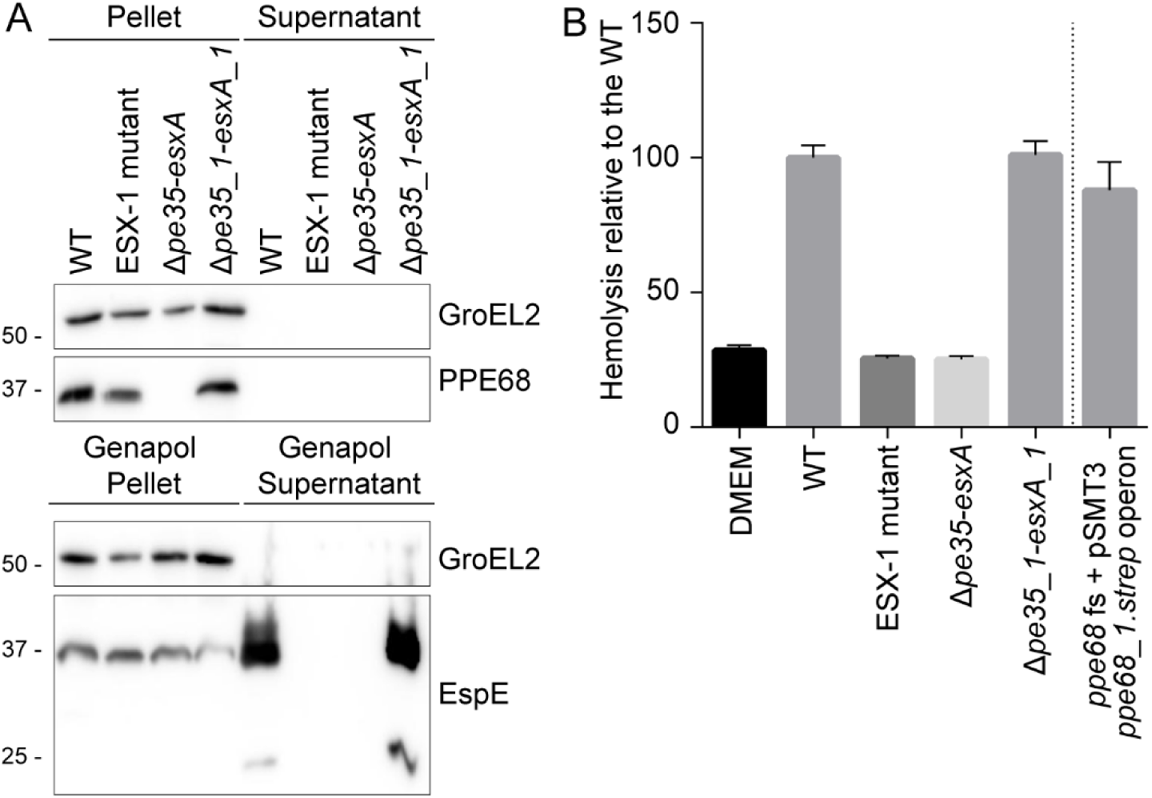
Deletion of the paralogous *ppe68_1* operon does not affect EspE secretion or hemolytic activity of *M. marinum*. A) SDS-PAGE and immunostaining was performed on the cell pellet, culture supernatant, Genapol pellet and Genapol supernatant fractions of *M. marinum* WT, an ESX-1 mutant (*eccC_b1_* mutant; M^VU^), the *ppe68* operon mutant (Δ*pe35-esxA*), and the *ppe68_1* operon mutant (Δ*pe35_1-esxA_1*). Proteins were visualized with anti-EspE and anti-PPE68 (ESX-1 substrates) antibodies. A processed band of 25 kDa was detected by the anti-EspE antibody in the Genapol supernatant fraction of the WT, which has been reported before (44). As a loading and lysis control, blots were incubated with antibodies directed against the cytosolic GroEL2 protein. In all blots, equivalent OD units were loaded: 0.2 OD for pellet and Genapol pellet and 0.4 OD for supernatant and Genapol supernatant fractions. B) Hemoglobin release after incubation of the same strains as under A) with sheep red blood cells. Hemoglobin release was quantified by determining the OD_450_ of the medium after incubation.

**Figure S4.**
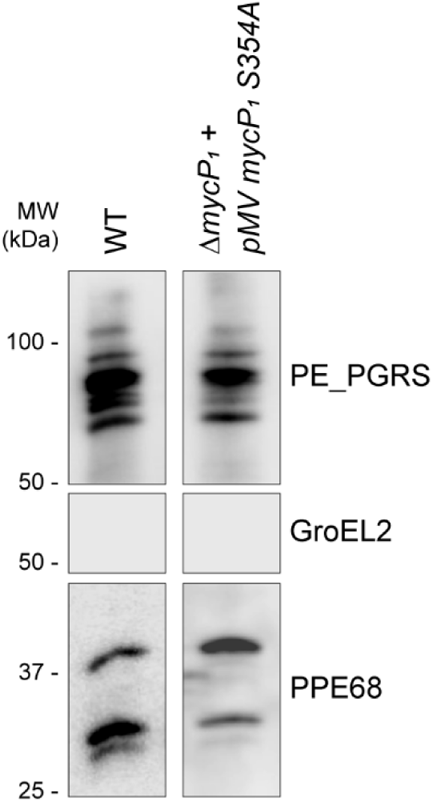
A point mutation in the active site of serine protease MycP_1_ does not affect the processing of PPE68 on the cell surface. SDS-PAGE and immunostaining of concentrated Genapol supernatant fractions of *M. marinum* WT and a *mycP_1_* deletion mutant complemented with a *mycP_1_* active site (S354A) variant, expressed from a pMV vector by the *hsp60* promotor. Proteins were visualized with an anti-PPE68 antibody. As a loading and lysis control, blots were incubated with antibodies directed against the cytosolic GroEL2 protein and the surface-localized PE_PGRS proteins. In all blots, equivalent amounts, i.e. 2 OD units, of Genapol supernatant fractions were loaded.

**Figure S5.**
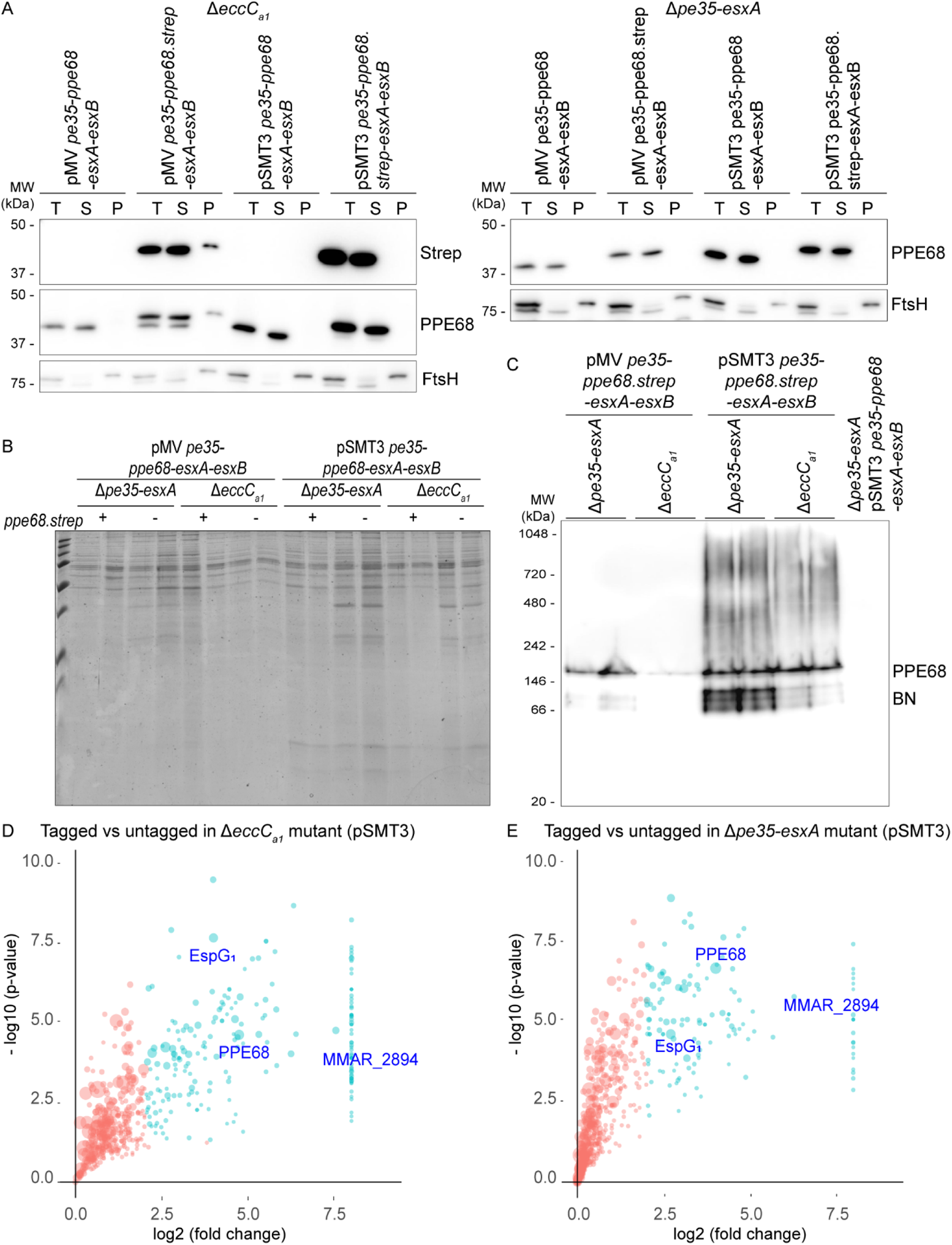
Purification of PPE68 from the *ppe68* operon mutant and the *eccC_a1_* mutant containing a pMV or pSMT3 vector with the *ppe68* operon of *M. marinum*. Genes were expressed from the *hsp60* promotor with or without a Strep-tag on the C-terminus of PPE68. A) Immunostaining of subcellular fractionations of the different strains. Proteins were visualized with anti-PPE68 and anti-StrepII antibodies, while anti-FtsH antibodies controlled for the loading and lysis. Notably, the FtsH antiserum cross-reacts with a cytosolic protein. In all blots, equivalent amounts corresponding to 0.2 OD units were loaded. T, total lysate, S, supernatant/soluble proteins, P, pellet/cell envelope proteins. B) Separation of the PPE68 purification samples by SDS-PAGE and subsequent staining with Coomassie Briliant Blue. C) Separation of complexes in the PPE68 purification samples by BN-PAGE and subsequent staining with Coomassie Briliant Blue. As a negative control, the purification sample of the *ppe68* operon mutant containing a pSMT3 vector with the *ppe68* operon, without C-terminal Strep-tag on PPE68, was included. D) Quantitative proteomics analysis comparing the PPE68 affinity purification from the soluble fraction of the *ΔeccC_a1_* mutant and the *ppe68* operon mutant (Δ*pe35-esxA*) producing PPE68 with or without a Strep-tag on the C-terminus, from a pSMT3 vector. Proteins that show log2 fold change > 2 and a −log10 p-value > 1.3 between tagged and untagged PPE68 are indicated as blue dots. Size of the dots are correlated to the number of spectral counts.

**Figure S6.**
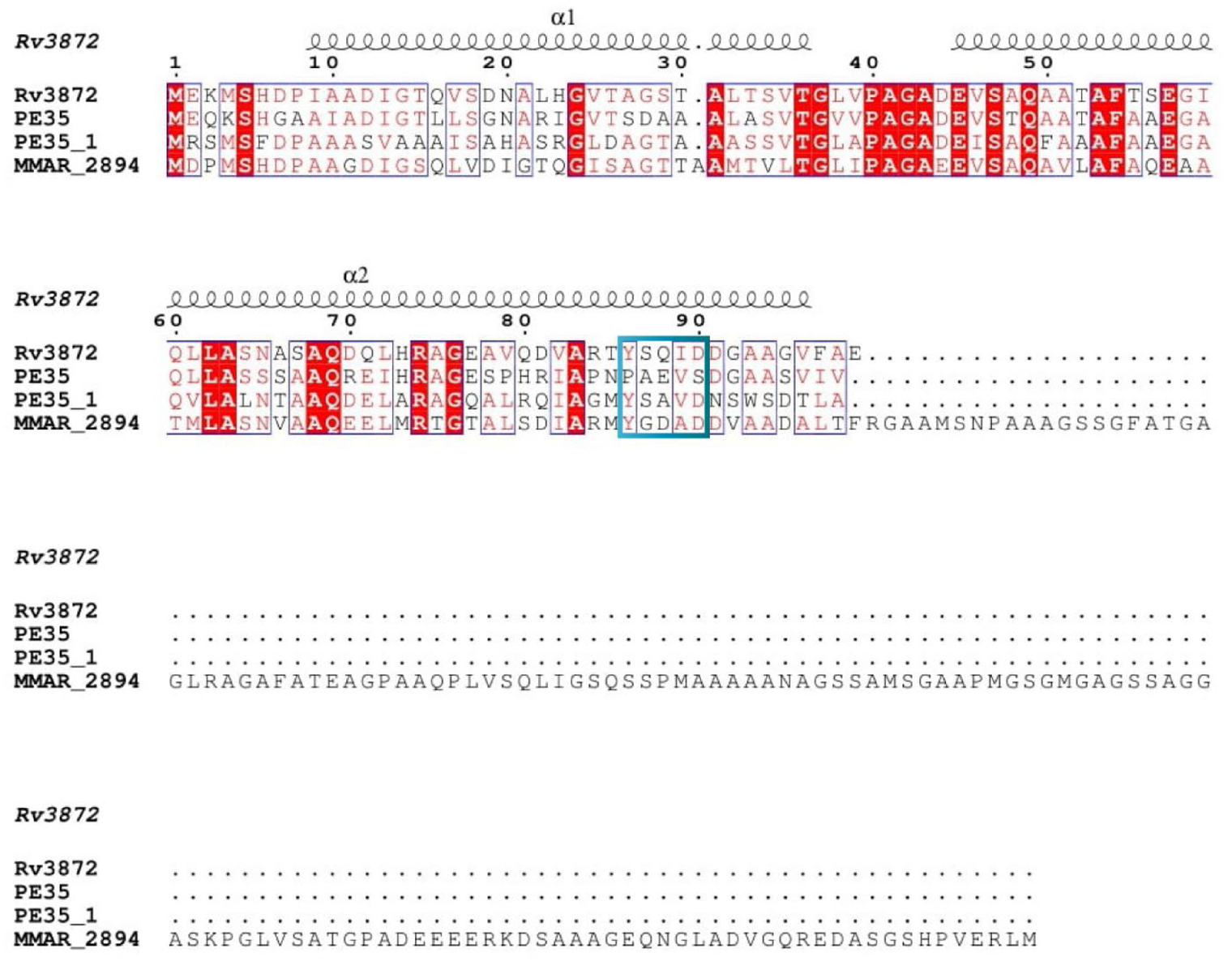
Protein sequence alignment of PPE35 homologs of *M. marinum* and *M. tuberculosis* including the secondary structure prediction of *M. tuberculosis* PE35. The secondary structure prediction was produced by Alphafold. The blue square indicates the conserved YxxxD/E secretion motif. Note that PE35 of *M. marinum* lacks this conserved motif. Rv3872 = PE35 of *M. tuberculosis*, PE35 = MMAR_5447 of *M. marinum* and PE35_1 = MMAR_0185 of *M. marinum*.

**Figure S7.**
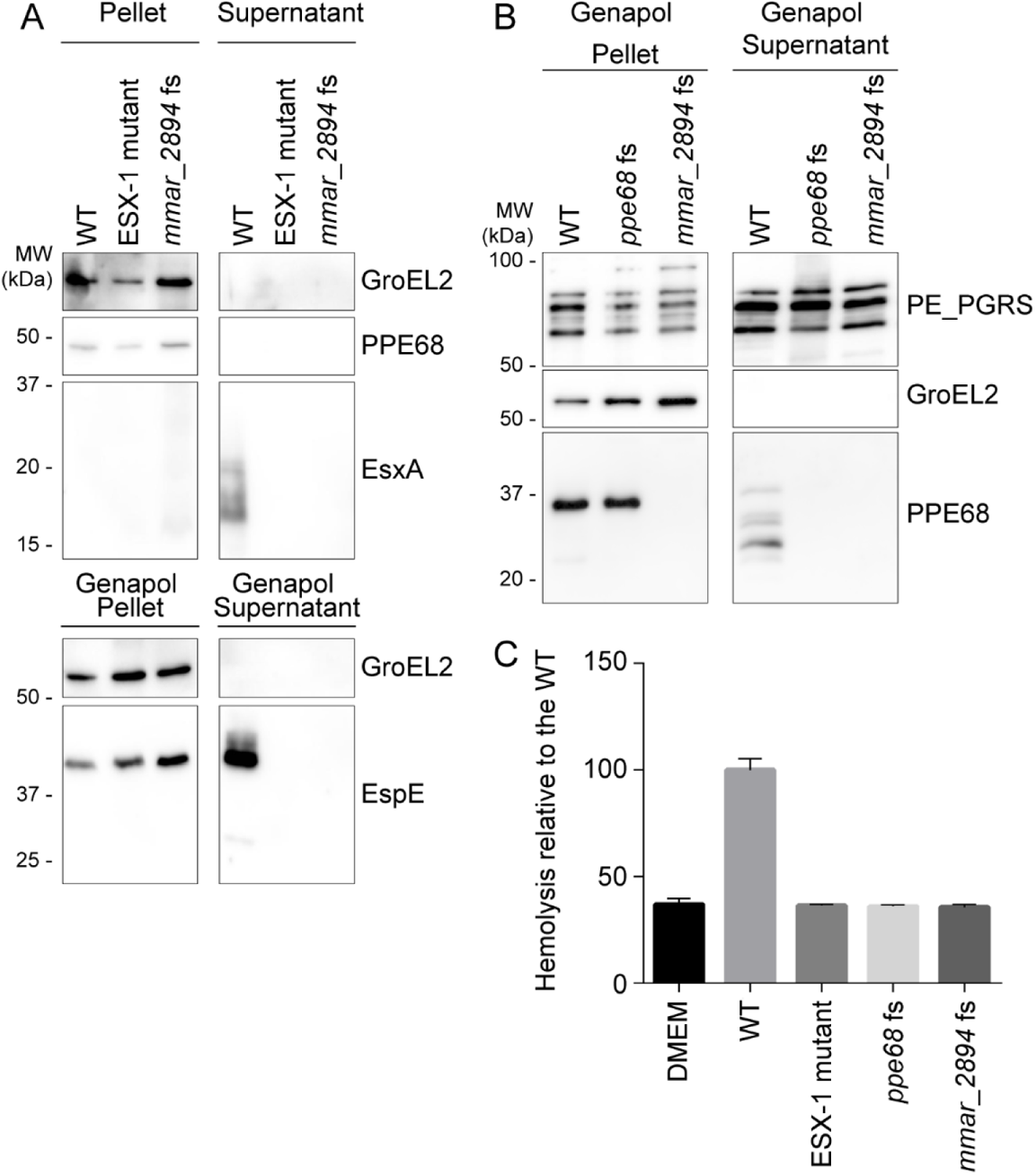
Secretion and hemolytic analysis of the *MMAR_2894* fs mutant. A) SDS-PAGE and immunostaining of the cell pellet, culture supernatant, Genapol pellet and Genapol supernatant fractions of *M. marinum*, WT, an ESX-1 mutant (*eccC_b1_* mutant; M^VU^) and the *MMAR_2894* fs mutant. Proteins were visualized with anti-EsxA, anti-EspE and anti-PPE68 antibodies (ESX-1 substrates). A processed band of 25 kDa was detected by the anti-EspE antibody in the Genapol supernatant fraction of the WT, which has been reported before (42). As a loading and lysis control, blots were incubated with antibodies directed against the cytosolic GroEL2 protein and the surface-localized PE_PGRS proteins. In all blots, equivalent OD units were loaded: 0.2 OD for pellet and Genapol pellet and 0.4 OD for supernatant and Genapol supernatant fractions. B) SDS-PAGE and immunostaining of the Genapol pellet and upscaled Genapol supernatant fractions of *M. marinum* WT, the *ppe68* fs mutant and the *mmar_2894* fs mutant. C) Hemoglobin release after incubation of the same strains explained under A and B with sheep red blood cells, quantified by determining the OD_450_ of the medium after incubation.

**Table S1.** Significant hits of PPE68 interactors as determined by protein purification and mass spectrometry analysis. Only proteins with an average normalized spectral counts >5 in the tagged samples, a log2 fold change > 2 and a −log10 p-value of > 1.3 between tagged and untagged samples are shown.

| *ΔEccCa1::pMV pe35-esxA* |  |  |  |  | *Δpe35-esxA::pMV pe35-esxA* |  |  |  |  |
| --- | --- | --- | --- | --- | --- | --- | --- | --- | --- |
| Protein | Average<br>raw<br>counts<br>tagged | Average<br>raw<br>counts<br>untagged | log2<br>fold<br>change | -log10<br>P-value | Protein | Average<br>raw<br>counts<br>tagged | Average<br>raw<br>counts<br>untagged | log2<br>fold<br>change | -log10<br>P-value |
| **EspG<sub>1</sub>** | 149.00 | 20.00 | 2.90 | 3.62 | **EspG<sub>1</sub>** | 150.50 | 17.50 | 3.10 | 5.99 |
| **PPE68** | 81.50 | 7.00 | 3.54 | 4.80 | **PPE68** | 115.00 | 8.50 | 3.76 | 3.58 |
| **MMAR_2894** | 15.50 | 0.00 | 8.00 | 2.59 | **MMAR_2894** | 36.00 | 0.00 | 8.00 | 5.79 |
|  |  |  |  |  | **NrdE** | 22.50 | 5.00 | 2.17 | 1.44 |
|  |  |  |  |  | **HemD** | 11.00 | 0.00 | 8.00 | 1.57 |
|  |  |  |  |  | **FadH** | 11.00 | 1.00 | 3.46 | 3.07 |
|  |  |  |  |  | **HupB** | 10.50 | 0.50 | 4.39 | 2.25 |
|  |  |  |  |  | **FtsE** | 9.00 | 1.50 | 2.58 | 2.02 |
|  |  |  |  |  | **MMAR_2279** | 8.50 | 0.00 | 8.00 | 1.34 |
|  |  |  |  |  | **FabD** | 8.00 | 0.50 | 4.00 | 1.59 |
|  |  |  |  |  | **RplW** | 8.00 | 1.50 | 2.42 | 2.11 |
|  |  |  |  |  | **RhlE** | 7.50 | 1.00 | 2.91 | 1.99 |
|  |  |  |  |  | **MMAR_5479** | 7.00 | 1.00 | 2.81 | 1.62 |
|  |  |  |  |  | **RpsS** | 6.50 | 0.00 | 8.00 | 1.59 |
|  |  |  |  |  | **PlsC** | 6.00 | 0.00 | 8.00 | 1.30 |
|  |  |  |  |  | **MMAR_1001** | 6.00 | 0.00 | 8.00 | 1.30 |
|  |  |  |  |  | **AhpC** | 5.50 | 0.00 | 8.00 | 2.70 |
|  |  |  |  |  | **RpmF** | 5.50 | 1.00 | 2.46 | 2.04 |
|  |  |  |  |  | **AsnB** | 5.50 | 1.00 | 2.46 | 1.54 |
|  |  |  |  |  | **ProB** | 5.50 | 1.00 | 2.46 | 1.44 |
|  |  |  |  |  | **ChoD_1** | 5.50 | 0.00 | 8.00 | 1.84 |

**Table S2.**
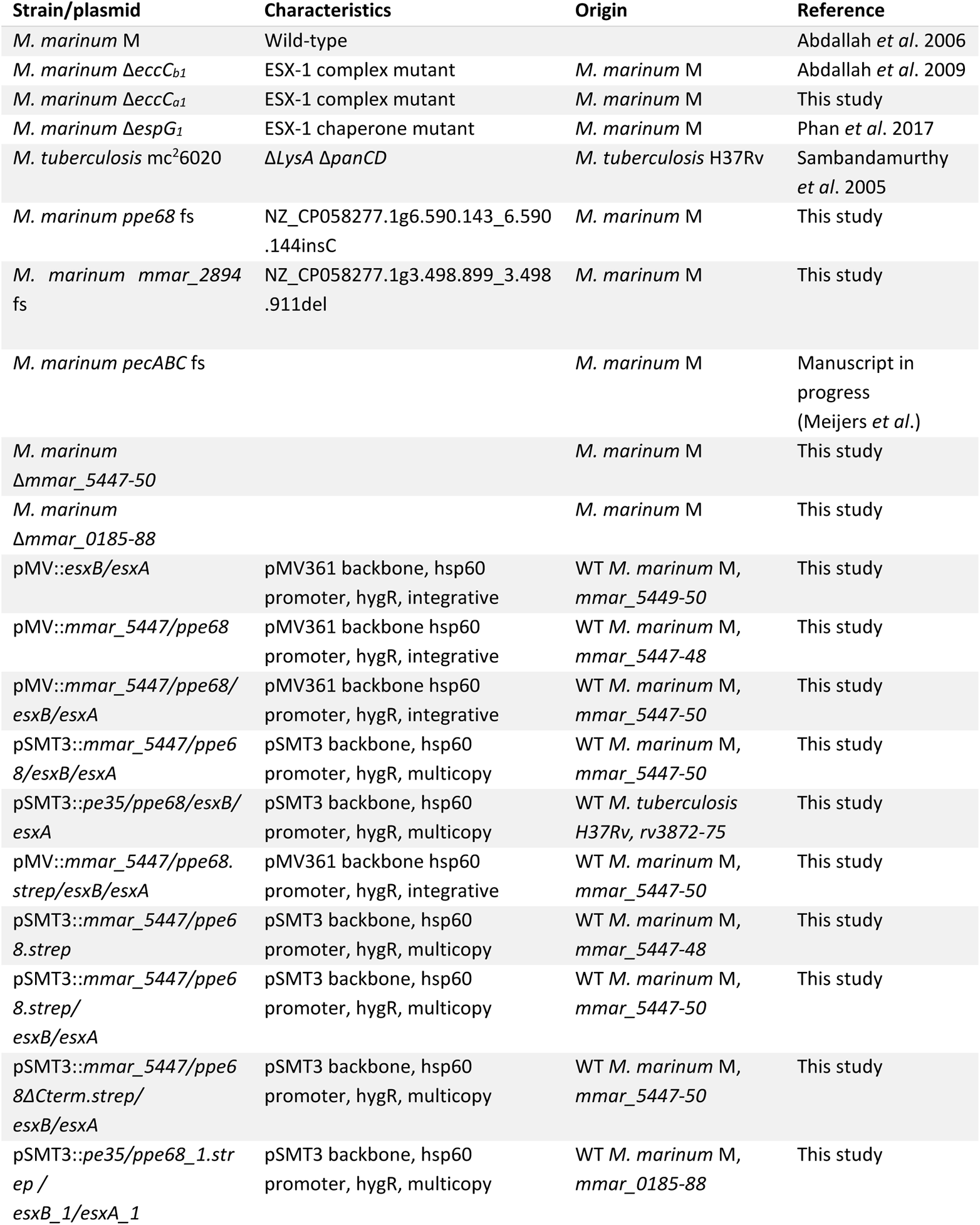
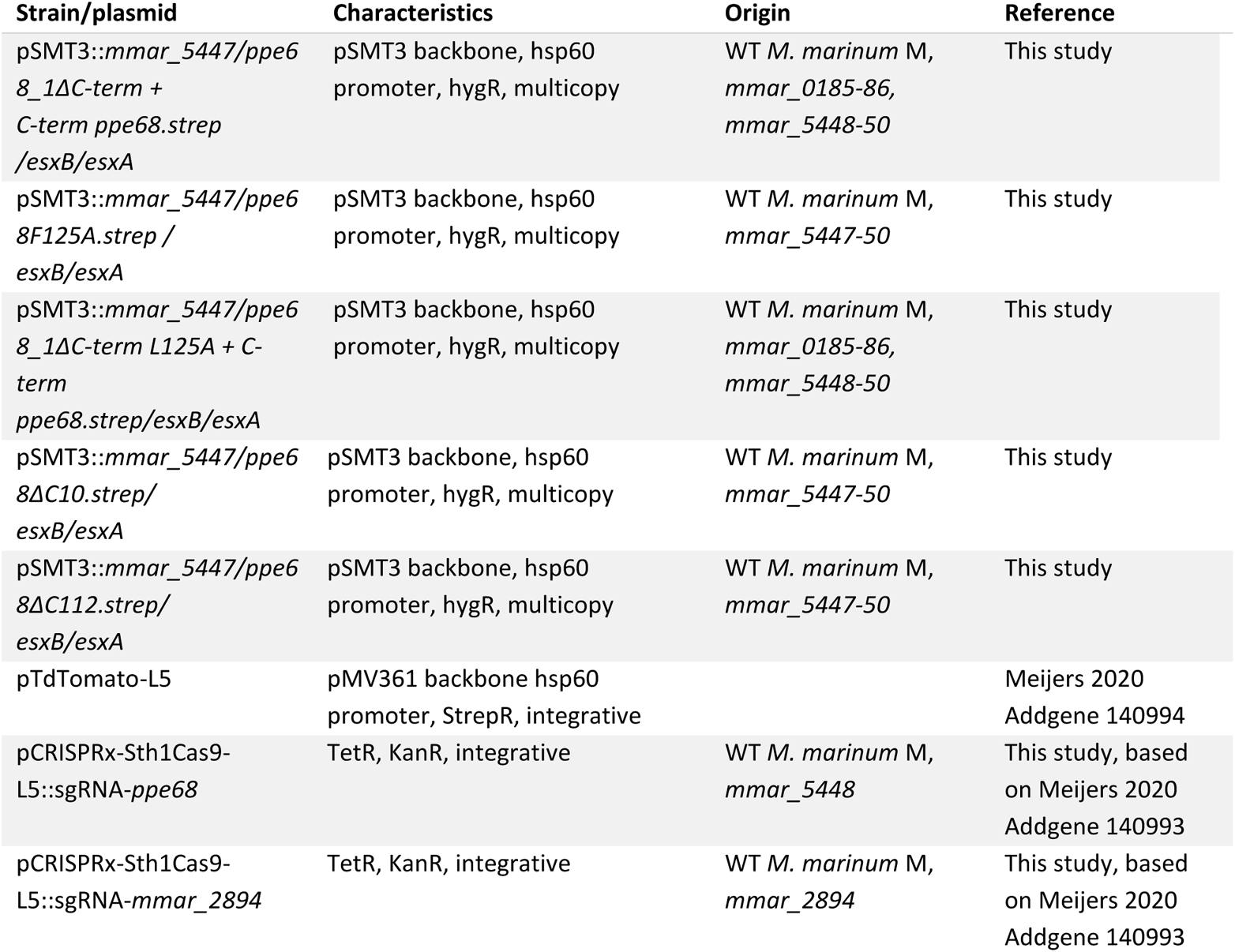
Strains and plasmids used in this study.

**Table S3.**
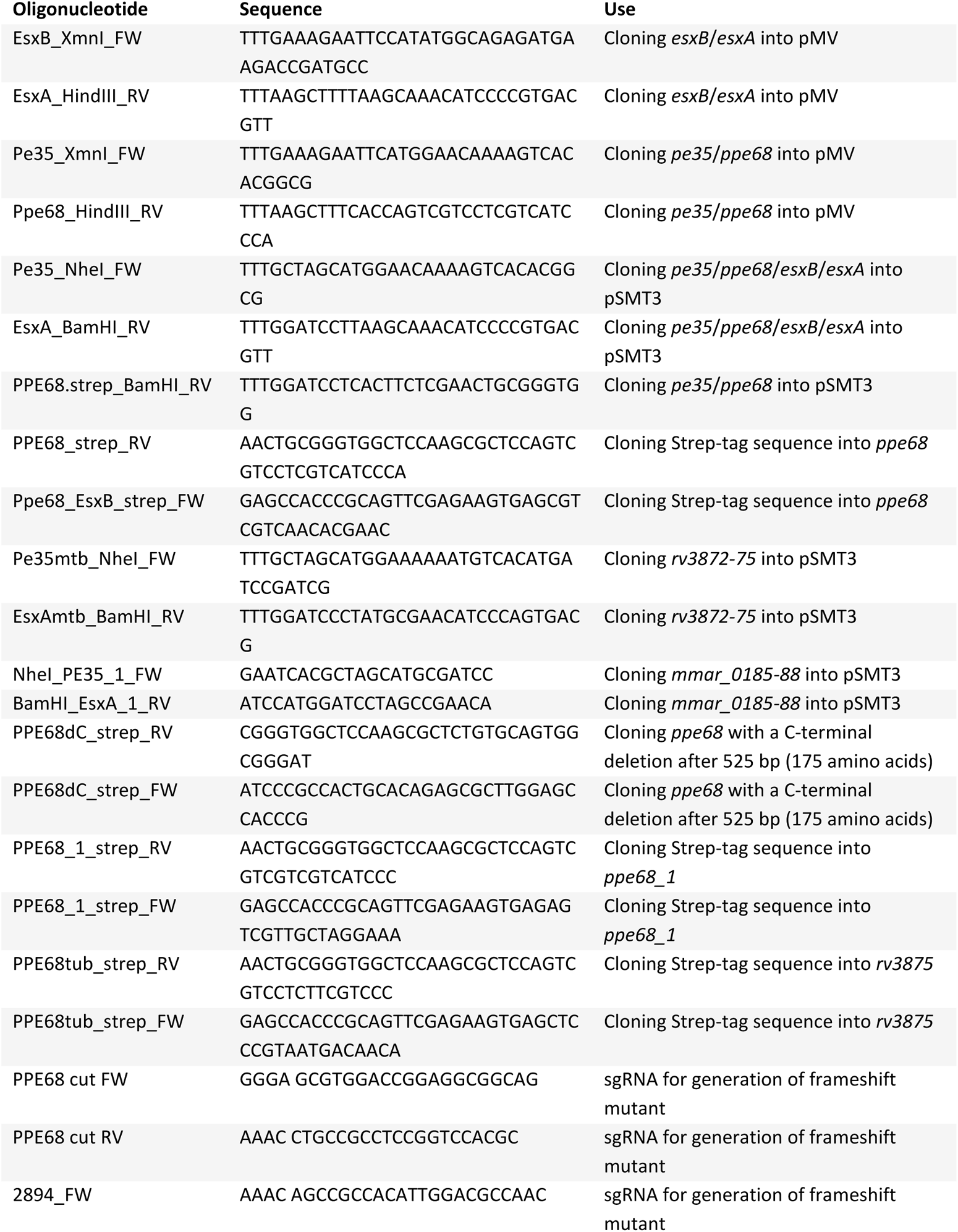
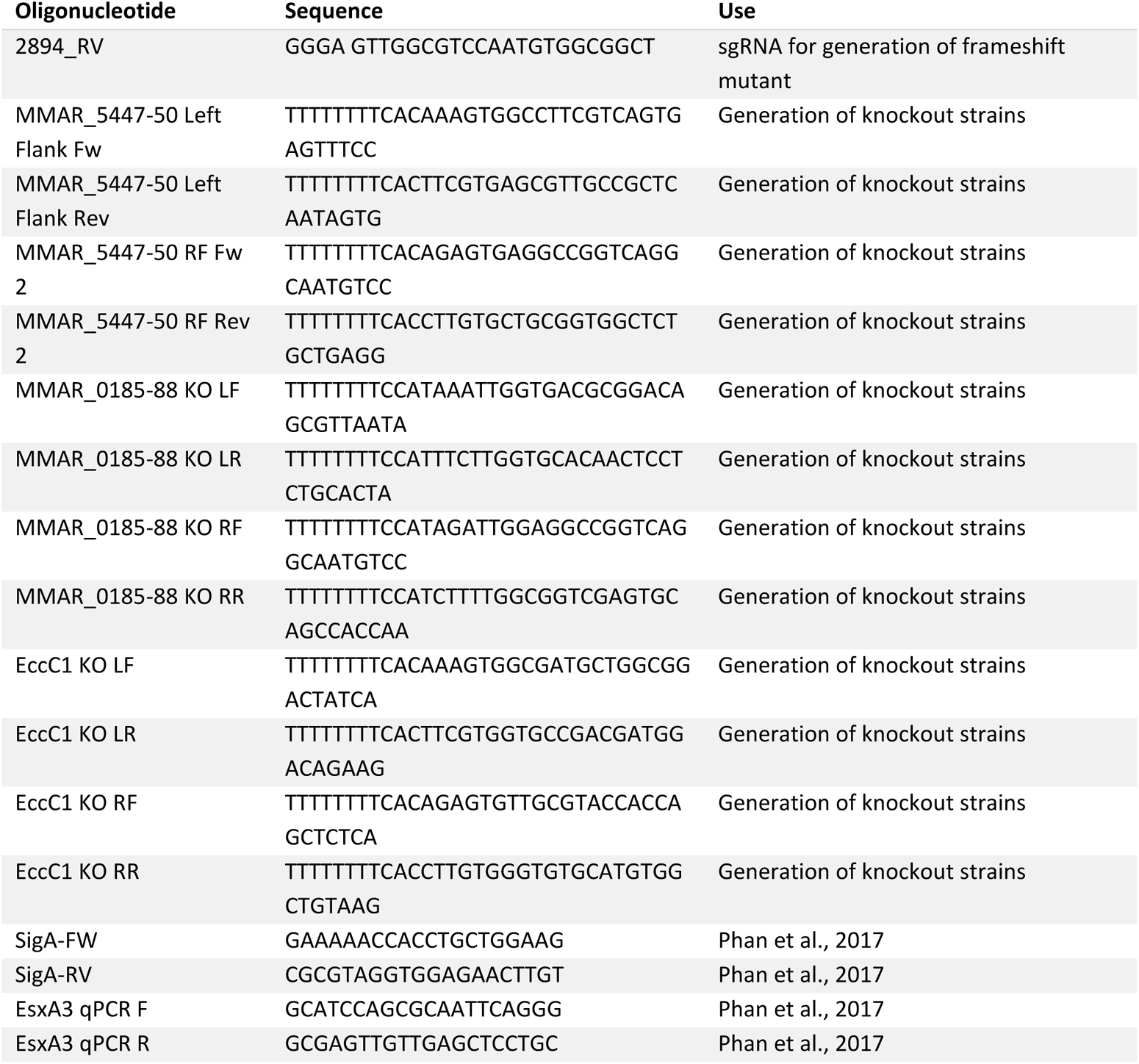
Primers used in this study.

